# Snowball Earths, population bottlenecks, and *Prochlorococcus* evolution

**DOI:** 10.1101/2020.11.24.395392

**Authors:** Hao Zhang, Ying Sun, Qinglu Zeng, Sean A. Crowe, Haiwei Luo

## Abstract

*Prochlorococcus* are the most abundant photosynthetic organisms in the modern ocean. A massive DNA loss event occurred in their early evolutionary history, leading to highly reduced genomes in nearly all lineages, as well as enhanced efficiency in both nutrient uptake and light absorption. The environmental landscape that shaped this ancient genome reduction, however, remained unknown. Through careful molecular clock analyses, we established that this *Prochlorococcus* genome reduction occurred during the Neoproterozoic Snowball Earth climate catastrophe. The lethally low temperature and exceedingly dim light during the Snowball Earth event would have inhibited *Prochlorococcus* growth and proliferation and caused severe population bottlenecks. These bottlenecks are recorded as an excess of deleterious mutations that accumulated across genomic regions in the descendant lineages. *Prochlorococcus* adaptation to extreme environmental conditions during Snowball Earth intervals can be inferred by tracing the evolutionary paths of genes that encode key metabolic potential. This metabolic potential includes modified lipopolysaccharide structure, strengthened peptidoglycan biosynthesis, the replacement of a sophisticated circadian clock with an hourglass-like mechanism that resets daily for dim light adaption, and the adoption of ammonia diffusion as an efficient membrane transporter-independent mode of nitrogen acquisition. In this way, the Neoproterozoic Snowball Earth event altered the physiological characters of *Prochlorococcus*, shaping their ecologically vital role as the most abundant primary producers in the modern oceans.

## Introduction

*Prochlorococcus* are the smallest and most abundant photosynthetic organisms on Earth (1). They are prevalent throughout the photic zone of the oligotrophic oceans between 40 °N and 40 °S (1), where they account for more than 40% of the biomass and contribute almost half of the net primary production (2). *Prochlorococcus* have diversified into two major phylogenetic groups with distinct ecology (ecotypes), with the high-light (HL) adapted monophyletic group imbedded in the low-light (LL) adapted paraphyletic group (3). The distinct ecotypes of *Prochlorococcus* evolved different pigments, light-harvesting systems, and phycobiliproteins, which allowed for efficient light absorption in the water column (4), and thus increased growth rates and primary production (5).

*Prochlorococcus* genomes have been shaped by stepwise streamlining, including a major genome reduction in their early evolution and a few minor modifications that followed (6, 7). Modern marine *Prochlorococcus* lineages, in particular those with small genomes, show very low ratios of nonsynonymous (*d_N_*) to synonymous (*d_S_*) nucleotide substitution rates, suggesting that natural selection is a highly efficient throttle on the accumulation of deleterious mutations (i.e., nonsynonymous mutations) in *Prochlorococcus* (6, 8). On long time scales, however, nucleotide substitutions at synonymous sites become saturated, invalidating the use of *d_N_*/*d_S_* to infer selection efficiency in deep time (9). Thus, an alternative approach focuses instead on different types of nonsynonymous substitutions leading to radical versus conservative changes in amino acid sequences, with the former more likely to be deleterious (10, 11). Excess radical mutations accumulate from random fixations of deleterious mutations by genetic drift (i.e., reduced efficiency of selection). Using this approach reveals that the major genome reduction in *Prochlorococcus* took place under reduced selection efficiency (9) and implies that the ancient population went through severe bottlenecks as the likely result of environmental catastrophe.

The environmental context underlying *Prochlorococcus* genome reduction remains unknown, however, and precise molecular dating is needed to link this important evolutionary event to its possible environmental drivers. By implementing comprehensive molecular clock analyses, we now link the early major genome reduction event of *Prochlorococcus* to the Neoproterozoic Snowball Earth events. These catastrophic disruptions to the Earth system would likely have challenged warm-water-loving photosynthetic *Prochlorococcus*, with strong potential to cause the population bottlenecks inferred from the genome sequences described above. *Prochlorococcus* survived this catastrophe through likely gains and losses of key metabolic functions reconstructed from the same genome sequences, which have a far-reaching impact on their success in today’s oceans. *Prochlorococcus* are thus vital “guardians of metabolism” (12), shepherding genes critical to the functioning of the biosphere across environmental catastrophes, including global glaciations.

## Results and Discussion

*Prochlorococcus* experienced a massive gene loss event on the ancestral branch leading to the last common ancestor (LCA) of clades HL, LLI, and LLII/III (6, 7, 13). This is confirmed by our analysis, which reconstructed 366 and 107 gene family losses and gains on this branch, respectively (Fig. 1A & S1). On the same ancestral branch, it was shown that *d*_R_/*d*_C_ is significantly elevated compared to the sister ancestral branch leading to the LCA of the LLIV (9), which was validated here (both sign test and paired *t*-test, *p* < 0.001; Fig. 1B). These results confirm that the major genome reduction event occurring on this branch was likely driven by genetic drift as a result of one or recurrent population bottlenecks (9). Given the global distribution and abundance of *Prochlorococcus*, and cyanobacteria more generally, such a bottleneck would likely require a global-scale event, like an environmental or climate catastrophe (e.g. meteorite impact, large igneous province emplacement, or glaciation).

**Fig. 1.**
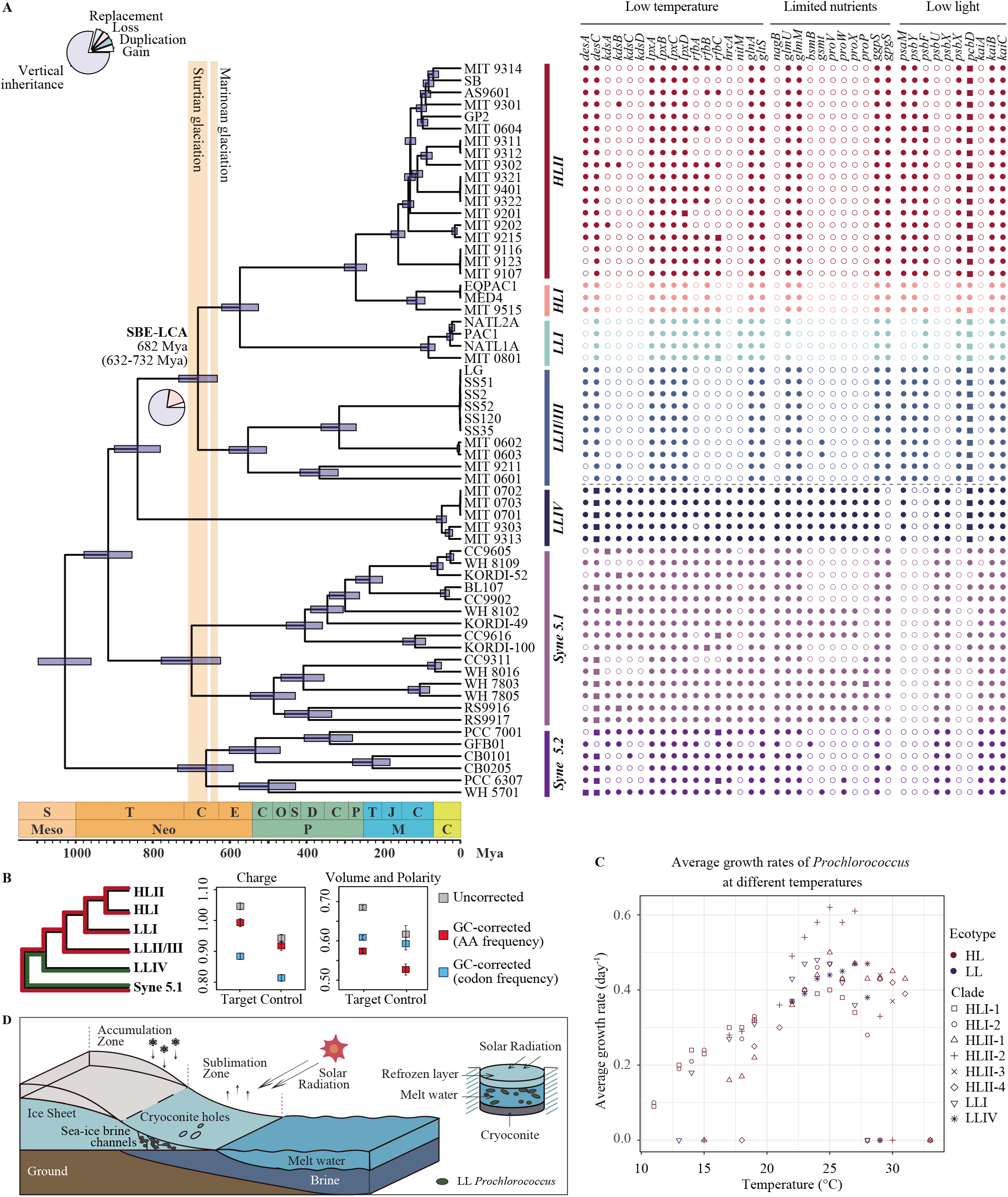
Evolution of *Prochlorococcus* during the Neoproterozoic Snowball Earth events. (A) (Left) Chronogram of the evolutionary history of *Prochlorococcus* estimated by MCMCTree. The evolutionary tree shown here is part of the species tree constructed with MrBayes based on 90 compositionally homogenous gene families shared by 159 cyanobacterial genomes (Fig. S4 D). Divergence time is estimated based on 27 gene sequences under calibration set C14 (Table S1). The vertical bars represent the estimated time of the Neoproterozoic glaciation events. The flanking horizontal blue bars on ancestral nodes represent the posterior 95% highest probability density (HPD) interval of the estimated divergence time. The pie chart on the ancestral branches leading to the node SBE-LCA provides the proportion of reconstructed genomic events including gene gain, gene loss, gene replacement, gene duplication and gene vertical inheritance. (Right) Phyletic pattern of key gene families that potentially enabled *Prochlorococcus* to survive harsh conditions during the Neoproterozoic Snowball Earth (at the ancestral node ‘SBE-LCA’). Solid square, solid circle and open circle next to each extant taxon represent multi-copy gene family, single-copy gene family, and absence of the gene family, respectively, in the genome. (B) (Left) The diagram helps understand how the *d*_R_/*d*_C_ was calculated. In this context, the ‘Target’ group includes all genomes of all HL clades, LLI and LLII/III, the ‘Control’ group includes all genomes of LLIV, and the ‘reference’ group includes all genomes of Syne 5.1. The *d*_R_/*d*_C_ for the ‘Target’ group (shown in Middle & Right) is calculated by comparing a genome from the ‘Target’ group to a genome from the ‘reference’ group (marked with red), followed by averaging the value across all possible genome pairs. Likewise, the *d*_R_/*d*_C_ for the ‘Control’ group (shown in Middle & Right) is calculated by comparing a genome from the ‘Control’ group to a genome from the ‘reference’ group (marked with green) and then by averaging the value across all possible genome pairs. (Middle & Right) The genome-wide means of *d*_R_/*d*_C_ values at the ancestral branch leading to SBE-LCA and that at its sister lineage. They were classified based on the physicochemical classification of the amino acids by charge or by volume and polarity, and were either GC-corrected by codon frequency (blue), GC-corrected by amino acid (AA) frequency (red) or uncorrected (gray). Error bars of *d*_R_/*d*_C_ values represent the standard error of the mean. (C) The average growth rate of *Prochlorococcus* ecotypes at different temperatures. Replicate cell cultures were grown in a 14:10 light: dark cycle at 66 ±1 μmol m^−2^s^−1^. The growth data used for plotting are collected from Johnson et al. 2006 and Zinser et al. 2007. (D) Diagram of putative bacterial refugia including cryoconite holes and sea ice brine channels in Neoproterozoic Snowball Earth.

To establish the environmental context for the large, ancient genome reduction, we estimated the timeline of *Prochlorococcus* evolution by implementing molecular clock analyses based on essential calibrations available in the cyanobacterial lineage. We recognize that the use of calibration sets adapted from previous studies (under calibrations C1-C8 in Table S1 with related references included there) results in up to ~320 Ma disparity (Fig. S2A) in the estimated time for the LCA of *Prochlorococcus* HL, LLI and LLII/III clades that emerged with the major genome reduction. We note that the calibrations in previous studies were not properly used. For example, the akinete fossil identified to 2,100 Mya was used as either the maximum bound or the minimum bound to calibrate the crown group of Nostocales (14, 15). However, given the fact that apomorphic character must evolve earlier than the divergence of crown group, morphological fossils can only serve as the minimum bounds on total groups of assigned lineages (16) (see Section 2.3 in Supplemental Methods for details). Thus, in the present study, we modified the calibration sets by constraining the lower bounds of the Nostocales (and the Pleurocapsales) total groups with morphological fossils and by leaving their upper bounds open (C9-C14; Table S1). Intriguingly, the variation is reduced to less than 10 Ma when these modified calibration sets are used (Fig. S2A).

Recent identification of non-oxygenic Cyanobacteria lineages such as Melainabacteria and Sericytochromatia as sister groups of oxygenic Cyanobacteria (17, 18) provides an alternative way to constrain the evolution of oxygenic Cyanobacteria. Specifically, given that oxygenic photosynthesis evolved at the stem lineage of oxygenic Cyanobacteria, we constrained the minimum age of the total Cyanobacteria group at 3.0 Ga, which is supported by geochemical evidence as the time when atmospheric oxygen became available (19, 20). To avoid the overly precise and potentially misleading age estimates, we calibrated the upper limit of the Cyanobacteria root using the ages when the planet Earth formed and became habitable (C15-C38 in Table S1; see Section 2.3 in Supplemental Methods for details). Using this strategy, we show that the age of *Prochlorococcus* major genome reduction remains stable when non-oxygenic Cyanobacteria outgroups were included (Fig. S2B). Since including the non-oxygenic Cyanobacteria have consistently reduced the precision of posterior age estimates, manifested as the higher slopes of the regression line between highest posterior density (HPD) width and the posterior age estimates compared to those without including these lineages (C15-C38 versus C1-C14 in Fig. S3; also see Section 2.6 in Supplemental Methods for an extended discussion), we focus on the crown oxygenic Cyanobacteria group dating (C7-C14) in the following discussions.

By comparing the width of the 95% HPD derived from each molecular clock analysis (Fig. S3), we inferred the most precise timeline of *Prochlorococcus* evolution (corresponding to the calibration set C14 in Table S1; see Section 2.3 in Supplemental Methods). Our time estimates revealed that the LCA of *Prochlorococcus* HL, LLI, and LLII/III clades diversified at 682 Mya (95% HPD 632-732 Mya), precisely dating the genome reduction event to this time. A 682 Mya date for the emergence of the LCA of *Prochlorococcus* HL, LLI, and LLII/III clades places the large genome reduction that took place in this lineage firmly within the Cryogenian Period (~720 to 635 Mya; Fig. 1A) and implicates the Snowball Earth icehouse climate conditions eponymous with the Period in the corresponding *Prochlorococcus* population bottleneck. We, therefore, refer to this ancestor as SBE-LCA (see Fig. 1A), short for “Snowball Earth” LCA. The Neoproterozoic climate catastrophe culminated in the Sturtian (~717 to 659 Mya) and Marinoan (~645 to 635 Mya) glaciations (Fig. 1A), which stretched from the poles to sea level near the equators, possibly wrapping the entire Earth under a frozen skin (21). This “Snowball Earth” persisted with the freezing temperature of seawater below the ice sheet lowered to −3.5°C (22). Since all assayed *Prochlorococcus* strains, including those affiliated with the basal LL ecotypes, reach maximum growth rates at approximately 25°C and rarely survive when the temperature drops to ~10°C (Fig. 1C) (23), we propose that extreme climate cooling during the Neoproterozoic Snowball Earth events was likely the major driver of severe bottlenecks in early *Prochlorococcus* populations.

Survival of *Prochlorococcus* populations through the Cryogenian would have required refugia, the nature of which would have shaped continued *Prochlorococcus* evolution. A variety of biotic refugia have been identified during Snowball Earth intervals, including the sea-ice brine channels within ice grounding-line crack systems (24) and cryoconite holes/ponds on the surface of the sublimation zone, which may have represented ~12% of the global sea glacier surface (25) (Fig. 1D). Despite providing the essential space for *Prochlorococcus* survival, these refugia would have presented a number of environmental stresses to *Prochlorococcus* populations such as low temperature, dim light, and limited nutrients (24–26). *Prochlorococcus* thus evolved a number of adaptive mechanisms to cope with these stresses via gene gains and losses, which we assessed by reconstructing the evolutionary paths of imprints that the Snowball Earth climate left in extant *Prochlorococcus* genomes.

Among these stresses, the most prominent was likely lethally low temperature. Maintaining membrane fluidity is of paramount importance under low-temperature conditions, which is largely achieved by the activities of fatty acid desaturase encoded by *desA* and *desC*. As a result, we inferred that these genes were retained in SBE-LCA (Fig. 1A). Lipopolysaccharide (LPS) in the outer membrane is known to provide the first line of defense against harsh environments (27), which contains the O-specific polysaccharide, the glycolipid anchor lipid A, and the polysaccharide core region. Based on our analyses, genes encoding the polysaccharide core region (*kdsABCD* for 3-deoxy-d-manno-octulosonate biosynthesis; Fig. 1A) were likely lost at SBE-LCA, while those encoding the other components were retained (*lpxABCD* and *rfbABC* for Lipid A precursor and O-specific LPS precursor biosynthesis; Fig. 1A). This inference is consistent with a previous conclusion that the loss of the LPS core region would increase the hydrophobicity and permeability of the cell envelope (28) to protect against cold conditions (29). Another metabolic modification in SBE-LCA was related to heat shock proteins (HSPs), which play crucial roles in tolerating environmental stresses including thermal shocks. Typically, HSPs are tightly regulated, as they respond quickly to stress and turn off rapidly once the stress disappears (30). However, the HSP repressor protein encoded by *hrcA* was inferred to be lost at SBE-LCA, which thus likely allowed the organism to continuously express HSPs to cope with prolonged lethally low temperature. In fact, constitutive expression of HSPs occurs in polar organisms such as the Antarctic ciliate *Euplotes focardii* (31) and the polar insect *Belgica antarctica* larvae (32). Extremely low temperature also made substrate acquisition difficult due to increased lipid stiffness and decreased efficiency and affinity of membrane transporters (33). Under such conditions, bacteria may increasingly rely on substrates whose uptake shows lower dependence on temperature. In sea-ice brines where less CO_2_ is dissolved (34), elevated pH promotes the conversion of ammonium to ammonia, which diffuses directly into cells without the aid of transporters in the membrane. Accordingly, species of bacteria and microalgae show a greater dependence on ammonium and ammonia at low temperatures and high pH than nitrate (35, 36), thereby reducing reliance on membrane transporters. In SBE-LCA, the potentially efficient utilization of ammonia made other N acquisition genes dispensable, leading to the neutral loss of nitrite transporter (*nitM*), whereas glutamine synthetase (*glnA*) and glutamate synthase (*gltS*) responsible for the utilization of ammonia after its assimilation were conserved (Fig. 1A).

An additional stress to *Prochlorococcus* during Snowball Earths was likely the extremely oligotrophic condition presented by bacteria refugia (37). Accordingly, SBE-LCA of *Prochlorococcus* evolved a few metabolic strategies for their survival. The amino sugar N-acetylglucosamine (GlcNAc) is used by bacteria such as *Corynebacterium glutamicum* as a carbon, energy, and nitrogen source (38). GlcNAc enters bacteria in the form of GlcNAc-6-phosphate (GlcNAc-6-P). However, instead of being metabolized, the loss of *nagB* for GlcNAc-6-P deamination at SBE-LCA suggests that GlcN6P is more likely to be involved in peptidoglycan (PG) recycling through the cascade catalysis by GlmM and GlmU (Fig. 1A) to generate UDP-GlcNAc, which is an essential precursor of cell wall PG and LPS (39). During cell turnovers, PG is continuously broken down and reused through the PG recycling pathway to produce new PG, and in some bacteria, PG recycling is critical for their long-term survival when growth is stalled under nutrient limitation (40). Thus, such a recycling mechanism seems to be key for the maintenance of cell integrity in SBE-LCA under oligotrophic conditions (25). Glycine betaine (GB) is known to be a ubiquitous protein-stabilizing osmolyte in bacteria, in particular cyanobacteria (41). However, genes involved in glycine betaine biosynthesis and transport were lost at SBE-LCA, including *bsmB* for dimethylglycine N-methyltransferase, *gsmt* for glycine/sarcosine N-methyltransferase, and *proVWXP* for glycine betaine/proline transport system (Fig. 1A). Instead, several other organic osmolytes might have been used during the Snowball Earths, as their biosynthetic genes were retained at SBE-LCA. The first examples are the *ggpS* gene encoding glucosylglycerol phosphate synthase for glucosylglycerol (GG) synthesis and the *gpgS* encoding glucosyl-phosphoglycerate synthase for glucosylglycerate (GGA) synthesis (Fig. 1A). Since the biosynthesis of GG and GGA requires less N compared to that of GB (42, 43), the potential use of GG/GGA instead of GB appeared favorable to SBE-LCA.

Low light intensity during Snowball Earth was another formidable challenge to phototrophs including *Prochlorococcus*. In the Neoproterozoic, the Sun was still at least 6% dimmer than that at present (44). Moreover, sea ice, especially when covered with snow, is an effective barrier to light transmission (45). This is in analogy to the deeper layers of today’s polar snow and glacier ice where irradiation is reduced and photosynthetic organisms and activities are scarcely detectable (46). *Prochlorococcus* may also need to compete with contemporary eukaryotic algae for the limited amount of light, because the latter became increasingly abundant during the Cryogenian glaciations and likely shared the same refugia with bacteria (37, 47). Consequently, photosynthetic organisms trapped in bacterial refugia or inhabiting waters below ice need to be physiologically geared to cope with low light. It was proposed that modification of the photosystem structure enables adaptation to the low light condition (48). We inferred a few changes in photosystem I and II (PSI/PSII) that occurred in SBE-LCA, including the gain of RC1 subunit PsaM, RC2 subunit PsbY, and an extra copy of the RC2 subunit PsbF, the loss of RC2 protein PsbU, and the replacement of RC2 subunit PsbX (Fig. 1A), but the molecular mechanism of these changes underlying low light adaptation is poorly understood. We also inferred an expansion of the *Prochlorococcus* antenna Pcb from two to six copies during the Snowball Earth (Fig. 1A), which may boost the light-harvesting capacity under low-light conditions (49).

Many cyanobacteria have a sophisticated circadian clock, which is essential in controlling the global diel transcriptional activities of the cells. This circadian oscillator system requires only three components: KaiA, KaiB, and KaiC (50). While all marine *Synechococcus* possess the three *kai* genes, most *Prochlorococcus* lack *kaiA* and, as a consequence, their circadian clocks rather behave like an “hourglass” which is reset every morning (51–53). Our analysis indicated that *kaiA* was lost at SBE-LCA (Fig. 1A). This is likely due to the prolonged darkness or low light conditions during the Snowball Earth, rendering the sophisticated circadian clock dispensable.

We argue that the genome reduction and metabolic adaptation events discussed above not only enabled *Prochlorococcus* to survive the Snowball Earth climate catastrophe, but also shaped the physiological characters and the biogeographic distribution of their descendants in the modern ocean. For example, the genome reduction that occurred in the early evolution of *Prochlorococcus* likely resulted in the reduced cell size and increased surface-to-volume ratio in their descendants, which may have enhanced their efficiency in nutrient acquisition (54) and eventually led them to dominate the photosynthetic communities in the most oligotrophic regions of today’s oceans (2). Likewise, new metabolic strategies that *Prochlorococcus* evolved to overcome the nutrient stresses during Snowball Earth, such as the recycling of cell wall components and the use of GG and GGA instead of nitrogen-rich GB as the organic osmolytes, decreased the nutrient requirements of the descendants’ cells and thus contributed to their success in the modern oligotrophic nitrogen-limited oceans. On the other hand, modifications of some important metabolic pathways may also have imposed deleterious effects on *Prochlorococcus* descendants. For example, whereas the replacement of circadian clock with an hourglass-like mechanism might have facilitated the ancestral lineage to adapt to the prolonged dim light condition during the Snowball Earth catastrophe, it likely prevents the dispersal of *Prochlorococcus* to high latitude regions in the modern ocean, where the day length varies substantially across seasons. Normally, organisms with circadian rhythms deal with these changes by anticipating the changes of light intensity and promptly regulating cellular processes such as DNA transcription and recombination via chromosome compaction, a known mechanism to protect DNA from UV radiation (55, 56). In the absence of the circadian clock, however, species such as *Prochlorococcus* cannot synchronize the endogenous oscillation with the environmental cycles and thus are at high risks of cell damages (57).

## Caveats and Concluding Remarks

The relaxed clock model implemented in the present study takes into account the rate variation among species and allows us to estimate a reliable timeline of *Prochlorococcus* evolution. On this basis, we link an ancestral phylogenetic branch that supported population bottlenecks and genome reduction of *Prochlorococcus* to the Neoproterozoic Snowball Earth. Using ancestral gene gain and loss analysis, we further identify potentially important metabolic strategies that the Cryogenian *Prochlorococcus* evolved to survive the glacial catastrophe. Despite these fascinating results, there are important caveats. We postulated that the icehouse conditions during the Cryogenian were lethal to the ancestral *Prochlorococcus* and thus inducing population bottlenecks. Apparently, this key assumption derives from our knowledge on the modern *Prochlorococcus* populations which do not grow below ~10°C, but this physiological character is not necessarily transmissible to the ancestral population. We also postulated that the metabolic traits we discussed earlier, which allowed *Prochlorococcus* to survive the lethally low temperature, extremely oligotrophic condition, and low light intensity, each must have conferred a strong fitness advantage (i.e., large selection coefficient *s*) to the Cryogenian *Prochlorococcus* population. This is because for populations under powerful genetic drift and thus having small effective population sizes (*Ne*), which is the case for the Cryogenian *Prochlorococcus* populations, only those mutants that confer sufficiently large benefits (*s* > 1/*Ne*) can be promoted by positive selection (58). While the metabolic traits we discussed fit the geochemical conditions well, they need additional evidence to support the hypothesis that they were subjected to positive selection and facilitated *Prochlorococcus* adaptation in those harsh conditions. From the perspective of the dating methodology, the uncertainty of our analysis largely comes from the use of calibrations. Molecular clock analysis requires at least one maximum age constraint (59). However, available cyanobacterial fossils can only serve as the minimum bounds (16), and therefore our analysis has to rely on the maximum bound provided by the root. We took a conservative approach by successively increasing the maximum bound of the root from 3,800 Ma to 4,500 Ma, which showed that the posterior age of the ancestral node SBE-LCA increased only slightly by ~7% (Fig. S2 and Table S1) and thus strengthened our conclusion of the coincidence between SBE-LCA and Cryogenian Snowball Earth.

The Neoproterozoic Snowball Earth hypothesis was proposed decades ago, which claimed the entire extinction of the photosynthetic organisms (21). In contrast to the original “hard” version of the hypothesis, a modified “soft” version of the Snowball Earth hypothesis was later proposed to include the likely persistence of refugia across the Cryogenian Period, which allowed for the survival of bacterial and simple eukaryotic lineages (60, 61). Survivors of the Snowball Earth included photosynthetic microorganisms (62, 63), which enabled continuous primary production across the interval (60, 64). Like other autotrophic organisms at the base of a food web, the survival of *Prochlorococcus* was likely important in sustaining primary production, heterotrophy, and carbon cycling, as well as broader ecosystem functioning during the Snowball Earth glaciations (60). On the other hand, the population bottlenecks we learned from *Prochlorococcus* may not necessarily have occurred in other cooccurring photoautotrophic lineages. Take *Synechococcus*, which evolutionarily most closely related to *Prochlorococcus*, as an example. The modern *Synechococcus* have wider geographical distribution than *Prochlorococcus*, and those inhabiting higher latitude regions are known to be more adaptive to the lower seawater temperature (65, 66). Moreover, *Synechococcus* harbor more diverse pigments than *Prochlorococcus*, which allow them to live with a wider range of light niches (67). These unique traits may increase the survivorship of *Synechococcus* during the Snowball Earths and thus reduce the chance to detect population bottlenecks, if any, in that difficult time.

On the other extreme, a few studies have proposed that microbial communities might have been only mildly affected by the Snowball Earth climate catastrophe (63, 64). These inferences were based on the microfossil and biomarker records, which, due to the lack of lineage-specificity, did not capture the nuances required to reconstruct effects on many ecologically important lineages as the *Prochlorococcus* studied here. Instead, we find that substantial disruptions to the Earth system, like the Neoproterozoic Snowball Earth, leave indelible signatures in microbial genomes, such that these heritable changes allow us to reconstruct interactions between environmental change and biological evolution deep in Earth’s history. By employing the accelerated genome-wide accumulation of the deleterious type mutations as a proxy for a rapid decrease in the population size of ancient lineages, we uncovered severe bottlenecks that shaped the early evolution of *Prochlorococcus* lineages. The precise molecular clock analyses as well as the ancestral genome reconstruction, furthermore, enabled us to link dynamics in ancestral population sizes to changes in metabolic potential and adaptation to icehouse climates through natural selection. Collectively, our findings demonstrate how paleomicrontological approaches can be used to connect large-scale dynamics in the Earth System to the genomic imprints left on extant microorganisms, which shape their ecological role and biogeographic distribution in the world today. They also illustrate how *Prochlorococcus* acted as important “guardians of metabolism” (12), safeguarding photosynthetic metabolic potential across the Snowball Earth climate catastrophe.

## Materials and Methods

Genomic sequences of Cyanobacteria were downloaded from public databases and manually annotated (see “Dataset_1.tbl” in online GitHub repository and Section 1 in Supplemental Methods). Divergence time of *Prochlorococcus* was estimated with MCMCTree v4.9e (68) on top of 27 genes (see “Dataset_2.tbl” in online GitHub repository) previously proposed to be valuable to date bacterial divergence (69) and *Cyanobacteria* phylogenomic trees. In previous studies, the LPP (*Leptolyngbya*, *Plectonema*, and *Phormidium*) group of *Cyanobacteria* located either at the basal of the Microcyanobacteria group (14, 70) or at the basal of the Macrocyanobacteria group (71, 72). Our analysis showed that this controversy is likely caused by the inclusion of composition-heterogeneous proteins, and that using composition-homogeneous proteins led to consistent support for the former hypothesis (Fig. S4; see “Dataset_3.tbl” in online GitHub repository and Section 2.2 in Supplemental Methods). Since molecular dating analysis is known to be intrinsically associated with calibration points (73), we summarized the calibrations of Cyanobacteria used in previous studies and modified them for our analyses with caution. Moreover, we proposed a new strategy to use calibrations when non-oxygenic Cyanobacteria were used as outgroups (Table S1; see Section 2.3 in Supplemental Methods for justification). We further assessed the fitness of different molecular clock models implemented in MCMCTree by using the package “mcmc3r” v0.3.2, based on which we decided to use the independent rates model for further molecular clock analyses. For each molecular clock analysis, the software ran twice with a burn-in of 50,000 and a total of 500,000 generations. The convergence was assessed based on the correlations of posterior mean time of all ancestral nodes between independent runs (Fig. S5). By implementing statistical tests based on the “infinite-site” theory (Fig. S3; see Section 2.6 in Supplemental Methods) we were able to select the most precise estimates of *Prochlorococcus* evolutionary timeline for illustration (Fig. S6) and further discussion.

Evolution of genome content via gene gains and losses was inferred using two independent methods, AnGST (74) and BadiRate v1.35 (75). The former assumes that the statistically supported topological differences between a gene tree and the species tree result from evolutionary events (gene loss, gene duplication, HGT, gene birth, and speciation), and infers these evolutionary events by reconciling the topological incongruences under a generalized parsimony framework by achieving a minimum number of the evolutionary events along the species tree, with penalties of an evolutionary event determined by the genome flux analysis (74). The latter does not rely on the tree topological incongruence information, but instead uses a full maximum-likelihood approach to determine the gene family turnover rates that maximize the probability of observing the gene count patterns provided by the family size table. The BadiRate analyses were run using nine strategies each with a distinct turnover rate model and a distinct branch model. The likelihoods of different runs were compared, and three strategies with the highest likelihood values were used (Fig. S7A). Further, results derived from AnGST (Fig. S7B) and BadiRate were compared and summarized to determine the common patterns shared by the two software (Fig. S7C), and important functional genes discussed were consistently inferred by these two methods. As the two methods inferred the qualitatively same pattern of genome size reduction on the branches leading to SBE-LCA, the number of gene gains and losses derived from the AnGST analysis was presented.

The inference of a potential change of selection efficiency on a given branch was performed by comparing the genome-wide *d*_R_/*d*_C_ value across single-copy orthologous genes of the branch to that of the closest sister branch. The *d*_R_/*d*_C_ value was calculated using RCCalculator (http://www.geneorder.org/RCCalculator/; see Section 4 in Supplemental Methods) based on two independent amino acid classification schemes (Table S2).

## Data Availability

The custom scripts as well as the sequence datasets are available in the online GitHub repository (https://github.com/luolab-cuhk/Prochl-SBE).

## Acknowledgments

We thank Allison Coe, Erik Zinser, and Zackary Johnson for providing the data of *Prochlorococcus* growth rates, Sishuo Wang, Tianhua Liao and Xiaoyuan Feng for their suggestions on molecular dating analyses. This work is supported by the National Natural Science Foundation of China (92051113), the Hong Kong Research Grants Council General Research Fund (14110820), the Hong Kong Research Grants Council Area of Excellence Scheme (AoE/M-403/16), HKU FoS funds to SAC, and the Direct Grant of CUHK (4053257 and 3132809).

