## Supplementary Information for "Snowball Earths, population bottlenecks, and *Prochlorococcus* evolution"

Haiwei Luo

**This PDF file includes:**

Supplemental Methods

References

Figures S1 to S7

Tables S1 to S2

### Supplemental Methods

### Table of Contents


**1. Taxon sampling and gene annotation of Cyanobacteria genomes**

**2. Timing the evolution of *Prochlorococcus***

*2.1 Relaxed molecular clock method implemented in MCMCTree*

*2.2 Phylogenomic tree construction of Cyanobacteria*

Ortholog identification among oxygenic Cyanobacteria genomes

Phylogenomic tree construction of oxygenic Cyanobacteria based on the complete set of orthologs

Phylogenomic tree construction of oxygenic Cyanobacteria based on the compositionally homogeneous subset of orthologs

Phylogenomic tree construction of Cyanobacteria phylum based on the subset of orthologs showing compositionally homogeneity

Resolved phylogeny of Cyanobacteria

*2.3 Justification of calibrations used for the molecular dating analyses*

Calibration of the Nostocales group

Calibration of the Pleurocapsales group

Calibrations of the root of oxygenic Cyanobacteria

Calibrations of the root of Cyanobacteria phylum

*2.4 Selection of molecular clock model*

*2.5 Input sequence data for molecular clock analysis*

*2.6 Assessing the precisions of molecular clock analyses*

**3. Reconstruction of gene gain and loss processes**

*3.1 Using AnGST*

*3.2 Using BadiRate*

*3.3 Gene gain and loss data integration*

**4. Calculating the rate of nonsynonymous nucleotide substitutions leading to radical and conservative amino acid changes, respectively**

### 1. Taxon sampling and gene annotation of Cyanobacteria genomes

By the time of this study (Dec 2018), a total of 309 oxygenic cyanobacterial genomes were available in the NCBI RefSeq database (1), among which 126 were marked as high-quality reference or representative genomes. For refined phylogenomic and relaxed molecular clock analyses, the 126 reference or representative genomes in RefSeq and the *Prochlorococcus* and *Synechococcus* collection included in a previous study (2) were used here, with the total number of 159 genomes (see "Dataset_1.tbl" in online GitHub repository). The latter genomes were included here because they were used to demonstrate an evolutionary mechanism underlying genome reduction of *Prochlorococcus* (2), which forms the basis of the present study. Clusters of orthologous group (COG) assignments for protein sequences were performed using RPSBLAST against the NCBI COG database (Dec. 2014 release) (3). Only the top COG hit for each protein was retained, which satisfied the domain-specific score threshold compiled from NCBI-curated domains and an e-value cutoff of $1e^{-3}$. Additional functional annotations were carried out using the KEGG database (2017 release) by BLASTP v2.2.6 and subsystem annotations at the RAST Server platform (4).

### 2. Timing the evolution of *Prochlorococcus*

#### *2.1 Relaxed molecular clock method implemented in MCMCTree*

The molecular clock hypothesis provides a powerful way to estimate species divergence time based on molecular sequences (5). Based on this theory, the genetic distance of two homologous sequences increases linearly with the length of time since their separation (5). However, evolutionary rates are often not constant over time and among lineages, which renders the strict clock hypothesis problematic in deep lineages like the Cyanobacteria studied here, and only occasionally useful for trees with shallow roots (6, 7). For this reason, we employed the software MCMCTree (8) to perform relaxed molecular clock analysis, which is known to be intrinsically associated with the use of phylogenomic tree, fossil calibrations, clock model and input sequence data. All these factors have been discussed below.

#### *2.2 Phylogenomic tree construction of Cyanobacteria*

The prerequisite for a reliable estimate of divergence time is to have a resolved phylogeny (9). In the case of Cyanobacteria, the mainly unresolved part resides at the LPP lineage, which contains *Leptolyngbya*, *Plectonema*, *Phormidium*, and *Synechococcus* sp. PCC7335 (10). In published phylogenies, LPP was a monophyletic group either located at the basal of the Microcyanobacteria group which contains *Synechococcus* and *Prochlorococcus* (11, 12), or at the basal of the Macrocyanobacteria group which contains the $N_2$-fixing Pleurocapsales and Nostocales (13, 14). We intended to solve this phylogenetic discrepancy before performing the time estimation.

<u>Ortholog identification among oxygenic Cyanobacteria genomes</u>

Using these 159 oxygenic Cyanobacteria genomes, we identified 381 single-copy orthologous gene families present in at least 155 genomes by implementing the orthology matrix algorithm (OMA v2.1.1) (15). We further examined whether all the members of each family shared the same COG functional category, and screened for potential inter-phylum horizontal gene transfer (HGT) using a BLASTP-based protocol similar to the one used in a previous study (16). A potential inter-phylum HGT event was defined as a cyanobacterial query with a non-cyanobacterial top hit (excluding the query itself; with an e-value ≤ 1e-10 and a percent of identity ≥ 35%) from the NCBI nr database (17). Finally, a total of 214 (out of 381) single-copy orthologous gene families that met all the requirements were retained for downstream analyses (see "Dataset_3.tbl" in online GitHub repository).

<u>Phylogenomic tree construction of oxygenic Cyanobacteria based on the complete set of orthologs</u>

The orthologous protein sequences were aligned using the E-INS-I refinement method of MAFFT v7.271 (18), and gaps were removed. The concatenation of the 214 single-copy orthologous gene families resulted in an alignment of 65,818 amino acid sites. PartitionFinder v2.1.1 (19) was used to determine the optimal partitioning schemes and best-fitting models using a greedy search with Bayesian information criterion (BIC). Phylogenetic analyses were performed using RAxML v8.2.10 (100 bootstrap replicates with GAMMA model of rate heterogeneity applied to each partition) (20) (Fig. S4A) and MrBayes v3.2.6 (21) (Fig. S4C). Each Bayesian execution computed two independent runs with four chains, running for 4,000,000 generations with a burn-in fraction of 25% and a sampling frequency of 2,000. Convergence between runs and posterior probabilities of the estimates was determined using Tracer v1.6 (22).

<u>Phylogenomic tree construction of oxygenic Cyanobacteria based on the compositionally homogeneous subset of orthologs</u>

An evident variation in G+C content among lineages suggested putative compositional heterogeneity across taxa (23). To assess whether each orthologous gene family significantly departs from the assumption of homogeneity, we carried out the simulation-based test implemented in the P4 phylogenetic toolkit (24), following a previous procedure in the analysis of Alphaproteobacteria phylogeny (25). For each individual ortholog alignment, we inferred the optimized parameters for the best-fitting substitution model based on ProtTest analysis (26), and used the resulting maximum likelihood (ML) tree as the phylogram on which 1,000 replicates were simulated. The distribution of amino acid compositions in the simulated data was subsequently compared with that of the empirical data under the $\chi^2$ statistic. Eventually, a set of 90 (out of 214) single-copy orthologous gene families confirmed compositional homogeneity at the 0.05 significance level (see "Dataset_3.tbl" in online GitHub repository). Phylogenetic analyses were performed again using these 90 families in the same way as elucidated above using both ML (Fig. S4B) and Bayesian (Fig. S4D) approaches, except that the MrBayes runs were ensured with convergence at the 3,000,000$^{th}$ generation (instead of 4,000,000$^{th}$) when the average standard deviation of the split frequency reached as low as 0.002 (< 0.01).

Phylogenomic tree construction of the Cyanobacteria phylum based on the subset of orthologs showing compositionally homogeneity

To incorporate non-oxygenic Cyanobacteria as outgroups in our molecular clock analyses, we obtained eight metagenome-assembled genomes (MAGs) of Melainabacteria and Sericytochromatia from GenBank. All these MAGs are known to be closely related to oxygenic Cyanobacteria, and have been used as outgroups in a previous study (27). We predicted protein sequences of these MAGs using the software Prokka v1.12 (28), which were then combined into the protein sequence dataset of oxygenic Cyanobacteria for another round of ortholog identification using OMA v2.1.1 (15). To simplify the process of phylogenomic tree construction, we extracted the previously identified set of compositionally homogeneous orthologs without additional simulation tests. They were used to build the phylogenomic tree of Cyanobacteria phylum using the software IQ-Tree v2.0 with automatically assigned amino acid substitution model under 1,000 ultrafast bootstraps (Fig. S4E).

<u>Resolved phylogeny of Cyanobacteria</u>

Using a concatenation of the protein sequences of the complete set (n=214) of single-copy orthologous gene families shared by 159 high quality oxygenic cyanobacterial genomes which contain more LPP members, the LPP lineage forms a polyphyletic group separately located at the basal of both Microcyanobacteria and Macrocyanobacteria in the ML (Fig. S4A) and Bayesian trees (Fig. S4C). Interestingly, using a concatenation of protein sequences of the remaining composition-homogeneous gene families (n=90), the phylogenies with the ML (Fig. S4B) and Bayesian (Fig. S4D) methods became fully congruent in which the LPP lineage became a monophyletic group and located at the basal of the Microcyanobacteria group. This phylogenetic structure has been commonly used in recent studies of time estimates (10-12), and remains stable when non-oxygenic Cyanobacteria outgroups were incorporated (Fig. S4E). We therefore employed the phylogeny shown in Fig. S4D and S4E for molecular dating analyses.

#### *2.3 Justification of calibrations used for the molecular dating analyses*

Molecular dating analyses are proposed to be intrinsically tied to calibration points (9). In the case of Cyanobacteria, there are two major ways to calibrate their evolution depending on whether the non-oxygenic Cyanobacteria lineages are used or not. In both ways, three time constraints are commonly used to calibrate the evolution of Cyanobacteria, which target the origin of oxygenic Cyanobacteria, the origin of Nostocales, and the origin of Pleurocapsales (10,

12, 29). However, when non-oxygenic Cyanobacteria lineages are included, additional time constraints on the root of Cyanobacteria phylum are required.

Despite the rigorous considerations of Cyanobacteria time constraints in previous studies, we notice that the way how fossil calibrations were applied in some of those studies was not appropriate (C1-C6 in Table S1). Thus, we modified the commonly used calibration sets in the present study (C7-C14 in Table S1) and also proposed a new strategy to calibrate the evolution of Cyanobacteria when non-oxygenic Cyanobacteria lineages are included (C15-C38 in Table S1). Details were provided below.

<u>Calibration of the Nostocales group</u>

The time constraints for the crown group of Nostocales have been heavily debated. The maximum boundary of Nostocales was set at different ages in previous studies. First, it was inferred based on heterocysts, which are specialized cells for nitrogen fixation under oxic conditions (30). As heterocysts were proposed to originate at the time when the atmospheric oxygen became increasingly available at 2,450 Mya (31, 32), this age was once set as the maximum boundary of Nostocales. Second, it was inferred based on akinetes, which is another type of differentiated cell of Nostocales for survival under extreme environmental conditions (32). Since Nostocales is not the only group in Cyanobacteria that produce akinetes, the age (2,100 Ma) of the earliest known akinetes fossil discovered in West Africa was used as the maximum boundary of Nostocales (12, 32). Third, the Nostocales cells are featured with morphological characters including the presence of sheath (condensed part of the akinete coat) and large cell diameter (13). As ancestral state reconstruction indicates that these characters occurred before the presence of Nostocales, the maximum age of Nostocales was set to 1,900 Ma when microfossils with both sheath and large cell diameter first appeared (13, 33). In terms of the minimum boundary, since the previously mentioned akinete fossil identified at 2,100 Ma was later inferred to be affiliated with Nostocales, the minimum boundary of Nostocales was set to 2,100 Ma in previous study (32, 34). An alternative minimum age of this lineage was set to 1,600 Ma due to the discovery of the nostocalean akinetes fossil in McArthur Group, northern Australia (35). We noticed that the akinete fossil identified to 2,100 Ma was used as either the maximum boundary or the minimum boundary of Nostocales in different Cyanobacteria dating analyses (10, 12). Although being self-contradictory, we still employed this boundary in different calibration sets (Table S1) for the purpose of comparison.

We note that morphological fossils such as akinetes and heterocysts have been used as the maximum bound to calibrate the crown group of Nostocales in previous studies (12, 29). However, given the potentially large gap between the initial appearance of an apomorphic character and its first fossilization time (36), the placement of these fossils on crown group of Nostocales may overly constrain the age prior and lead to false precisions in time estimates. Given the fact that apomorphic characters must have evolved earlier than the divergence of the crown group of assigned lineage, a more secure way to use these morphological fossils is to constrain the minimum age on total groups (36). From this perspective, the use of the nostocalean akinete fossils as the minimum constraints in previous studies are also inappropriate, as they were placed on the crown group of Nostocales (10, 29). Given these considerations, in the present study, we employed these morphological fossils to calibrate the lower bounds of the Nostocales total group regardless of whether the non-oxygenic Cyanobacteria lineages were used or not, and left the upper limit of Nostocales group open to avoid overly precise age estimates (C9-C38 in Table S1).

Calibration of the Pleurocapsales group

The time constraints for the crown group of Pleurocapsales are also contentious. Members of Pleurocapsales have large cell diameters (13). Since this character has been proposed to evolve earlier than the ancestor of Pleurocapsales, the maximum age of Pleurocapsales was once set to 2,450 Ma when the large cell diameter appeared in microfossil (13). Alternatively, since Pleurocapsales evolved later than filamentous and coccoid Cyanobacteria (12), which were proposed to occur at 1,900 Ma based on the microfossil identified in Gunflint chert (37), the maximum boundary of Pleurocapsales was set to 1,900 Ma in previous Cyanobacteria dating analyses (12, 29). The minimum age of Pleurocapsales was set to 1,700 Ma because of the microfossil identified in Hebei, China (38, 39).

We argue that the use of morphological fossils such as filamentous and coccoid cells as the maximum bound of Pleurocapsales in previous studies was not appropriate for the same reason we provided in the last section 'Calibration of the Nostocales group'. Thus, we modified the use of microfossils at 1,900 Ma as the minimum bound of total Pleurocapsales group. Moreover, since the maximum bound is hard to be established using fossil records (36), we left the upper limit of Pleurocapsales group open (C9-C38 in Table S1).

Calibrations of the root of oxygenic Cyanobacteria

For the root of oxygenic Cyanobacteria (i.e., the root of the phylogenomic tree when non-oxygenic Cyanobacteria lineages are not included; Fig. S4D), the minimum age at 2,320 Mya is commonly applied because of the convincing geochemical evidence for the rise of atmospheric oxygen at that time known as the Great Oxidation Event (GOE) (40), though recent studies showed that GOE may antedate the crown group of oxygenic Cyanobacteria (41, 42). The upper limit calibration of this root has been even more contentious. It was initially reported that 2-methylhopane can be used as a biomarker for Cyanobacteria (43), and the oldest record of this biomarker is dated back to 2,700 Mya (44), but the taxonomic link of 2-methylhopane to Cyanobacteria was challenged by the discoveries that 2-methylhopane is produced by the anoxygenic phototroph *Rhodopseudomonas palustris* under anaerobic conditions (45), and that the key gene for the methylation at the C-2 position of hopanoids was also found in α-Proteobacteria and Acidobacteria (46). The use of 2,700 Mya as the maximum age of the emergence of oxygenic Cyanobacteria was further weakened by a recent finding that the previously studied samples contained contaminants (47). On the other hand, ample geochemical evidence based on various sensitive redox proxies indicates the appreciable levels of the atmospheric oxygen at 3,000 Mya (48-50), which has been used as the upper bound of crown oxygenic Cyanobacteria in recent molecular clock analyses (12, 29). Consequently, in the cases when non-oxygenic Cyanobacteria lineages were not included, we calibrated the lower and upper limit of the crown oxygenic Cyanobacteria at 2,320 Mya and 3,000 Mya, respectively (C9-C14 in Table S1; Fig. S4D).

Recently, two lineages have been identified as the outgroups of oxygenic Cyanobacteria: Melainabacteria and Sericytochromatia (27, 51). Members of these outgroup lineages are proposed to lack essential genes for photosynthesis and carbon fixation, suggesting that the last common ancestor of Cyanobacteria was non-phototrophic (27). If this is the case, the oxygenic photosynthesis could be an evolutionary synapomorphy, which likely evolved at the stem lineage of oxygenic Cyanobacteria. Thus, when non-oxygenic Cyanobacteria lineages are incorporated, it is more appropriate to constrain the lower bound of total oxygenic Cyanobacteria instead of the upper bound of crown oxygenic Cyanobacteria using the geochemical evidence that atmospheric oxygen became available at 3,000 Mya (48-50) (C15-C38 in Table S1; Fig. S4E).

Calibrations of the root of Cyanobacteria phylum

In the cases when non-oxygenic Cyanobacteria lineages were included, we have to calibrate the root of phylogeny (i.e., the root of the Cyanobacteria phylum; Fig. S4E). To avoid overly precise age estimates, we constrained the upper limit of the Cyanobacteria root as ancient as possible. Given the potentially great influence of root prior on time estimates (52), we attempted different maximum prior ages for comparison. For example, we used 4,200 Mya, 4,000 Mya and 3,800 Mya by considering the time when the planet Earth became habitable and fostered the earliest life (53, 54) (C15-C32; Table S1). Additionally, a more conservative age at 4,500 Mya was used, since it was the time when the planet Earth formed (53) (C33-C38; Table S1).

*2.4 Selection of molecular clock model*

Molecular clock model is known to have a strong impact on posterior age estimates (55). The software MCMCTree implements different relaxed molecular clock models for time estimation, including auto-correlated rates (AR) model and independent rates (IR) model. The former assumes that the evolutionary rates in daughter species are statistically distributed around the parental rates, whereas the latter assumes a fully independent rate among evolutionary branches (56).

To assess the fitness of each clock model in our data, we compared the Bayes factors (BF) of these models using the thermodynamic integration method in the package “mcmc3r”

(56). While the method is powerful, it is very computationally intensive. Thus, we used the calibration set C9 as the representative for Bayesian model selection. Our results indicate that the IR model is superior to the AR model, as the BF value of the former is much higher than that of the latter (0.999 vs 0.001). We therefore employed the IR model in the following molecular clock analyses.

*2.5 Input sequence data for molecular clock analysis*

As an enlarged sequence dataset is able to improve the precision of time estimation based on the infinite-site theory (57), we employed as many as 25 core protein-coding genes (58) and two rRNA genes (16S, 23S) in the present study. Since substitutions at the third codon positions are largely silent and thus reach saturation rapidly, only the first and second codon positions of the 25 protein-coding genes were used. These 25 conserved protein-coding genes were previously identified from a genomic dataset spanning multiple bacterial and archaeal phyla and used to infer the evolutionary timeline of those groups (58). For each gene, we selected the best-fitting nucleotide substitution model by jModelTest (59), and calculated a rough substitution rate using BASEML (8) under a strict molecular clock. Further, the mean substitution rate was calculated based on the substitution rates of all input gene sequences, and then was used to inform the Dirichlet-gamma prior (rgene_gamma) in MCMCTree.

*2.6 Assessing the precisions of molecular clock analyses*

Evaluation of molecular clock analyses is important, since using different calibration set leads to a difference up to over 320 Ma in the estimates of the SBE-LCA when the non-oxygenic Cyanobacteria were not included (i.e., the last common ancestor of *Prochlorococcus* HL, LLI and LLII/III) (655 Mya under calibration set C6 vs. 981 Mya under calibration set C3; Fig. S2). Although the variation reduces to less than 10 Ma when the ages were estimated with the modified calibration sets (C7-C14 in Table S1), statistical evaluations of these analyses are valuable. The Bayesian inference approach that implemented in MCMCTree integrates the information from both calibrations and genetic data for posterior age estimation (60). Once the use of a calibration set is settled, according to the infinite-site theory, increased number of sites are recommended for molecular clock analysis, as they reduce the uncertainty in genetic distance estimate and increase the precision of the posterior time estimates (57). Theoretically, if sequences of infinite sites are used, the uncertainties in posterior time estimates are solely imposed by the uncertainties of the calibrations (57). By plotting the widths of 95% HPD interval against the posterior mean ages, we are able to assess the precision of the molecular clock analyses by comparing the slopes of the regression lines. A greater slope represents a lower precision of the time estimates (60, 61).

It has been repeatedly proposed that using multiple and more calibrations often lead to more reliable estimation than using less or even a single calibration (57, 62). Consistently, we showed that the time estimates based on calibration set C7 and C8 with a single calibration node has a high slope of 0.19 and 0.29, respectively (Fig. S3). It means that every 100 Ma divergence adds 19 and 29 Ma uncertainty in the posterior time estimates, respectively. According to this rule, the time estimates based on calibration set C14 has the lowest slope (0.149; Fig. S3), suggesting that the posterior time estimates of the SBE-LCA derived from this set are most precise. We did not further consider the analyses based on the calibration sets C1-C6 because the calibrations were not appropriately placed on the phylogeny.

We note that including Melainabacteria and Sericytochromatia consistently lead to less precise age estimates, as shown by higher slopes of the regression line between HPD width and the posterior age estimates (C15-C38 versus C1-C14 in Fig. S3). Given that genomes of these non-oxygenic Cyanobacteria lineages are fully represented by metagenome-assembled genomes (MAGs) but genomes of oxygenic Cyanobacteria used in our analyses are all derived from pure cultures, we hypothesize that the use of MAGs in molecular dating analysis, particularly those of lineages occupying important phylogenetic positions, may increase the uncertainties of the poterior age estimates. The quality of MAGs is questionable. While the CheckM (63) predicted that all of the MAGs used here show high level of completeness and low level of contamination (see "Dataset_1.tbl" in online GitHub repository), these assessments may not be reliable, as shown in a recent benchmarking study (64). For example, MAGs with estimated completeness as high as 95% may capture only three-fourths of the population core genes and a half of the variable genes, suggesting a greater amount of DNA is missing in the assemblies than estimated (64). Moreover, MAGs with estimated contamination as low as 1.5% may incorporate up to 5% of their genes with other taxonomic origins, suggesting a potentially higher contamination rate in the MAGs (64).

### 3. Reconstruction of gene gain and loss processes

#### *3.1 Using AnGST*

Genome content evolution via gene gains and losses was inferred using the gene tree vs. species tree reconciliation approach implemented in AnGST (65). The 62 genomes comprising the *Synechococcus-Prochlorococcus* monophyletic group were retrieved from the ultrametric cladograms yielded by the molecular dating analyses as described above (Fig. S7B). Gene trees were constructed using the following procedure. Firstly, homology relationships among proteins of the 62 *Synechococcus* and *Prochlorococcus* genomes were determined using OrthoFinder v2.2.1 (66) with DIAMOND as the alignment program (67). We identified 4,689 orthogroups (out of 5,615 orthogroups in total) each with at least three sequences. Next, multi-sequence alignments were constructed for each orthogroup using E-INS-I method implemented in the software MAFFT v7.222 (18), and trimmed with trimAl v1.4 ('-gappyout' option) to remove poorly aligned and excessively gapped regions (68). Lastly, gene trees were built using IQ-TREE v1.6.2 (69) under the ModelFinder feature (-m MFP) with ultrafast bootstrapping (1,000 replicates).

The reconciliation was inferred for each orthogroup under a generalized parsimony framework to achieve a minimum number of evolutionary events (gene loss, gene duplication, horizontal gene transfer [HGT], gene birth and speciation) along the species tree, using event penalties determined by the genome flux analysis (65). The genome flux analysis requires a minimal average difference in genome size between the ancestor and the descendant across the branches of the species tree, resulting in a set of optimized event penalties. We implemented the genome flux analysis with the speciation penalty fixed at 0.0 and the loss penalty at 1.0 as recommended in a previous study (65). The minimal genome flux was achieved when the HGT and duplication penalties are equal to 3.0 and 4.0, respectively). The HGT penalty inferred here agreed with the value achieved in a previous study based on a wide range of taxa across all three domains (65), and also confirmed HGT as the strongest effect on the genome flux as suggested in previous studies (65, 70, 71).

For all reconciliations performed, we enforced the time consistency (ultrametric = True) and restricted transfers to occur only between contemporaneous lineages. All 1,000 bootstrap replicates of each gene tree were provided to AnGST to resolve the gene tree phylogenetic uncertainties through amalgamation (65). AnGST incorporates the gene tree refinement procedure into the reconciliation process, and yields a chimeric gene tree (from the bootstrap replicates) which results in the lowest reconciliation cost, satisfying a generalized parsimony criterion (65). The numbers of gain, loss, and transfer events were summarized based on the AnGST output for each orthogroup across all branches along the species tree.

*3.2 Using BadiRate*

Gene gains and losses were also inferred with the likelihood-based method equipped in BadiRate v1.35 (72), which uses a full ML approach to determine the gene family turnover rates that maximize the probability of observing the gene count patterns provided in the family size table. A table of gene counts, consisting of all the aforementioned 5,615 orthogroups inferred by OrthoFinder v2.2.1 (66), and the same ultrametric time tree used in the AnGST analysis were used as the inputs. We fit nine different combinations of turnover rates (e.g., Birth-Death-Innovation model [BDI], Gain-Death model [GD], Lambda model [L] and Lambda-Innovation model [LI]) and branch models (e.g., Global-Rates model [GR], Branch-Specific-Rates model [BR], or Free-Rates model [FR]), including BDI/GD/L/LI+GR+ML, BDI/GD/L/LI+BR+ML and GD+FR+ML. Due to the computational intensity of the FR branch model, it was only implemented with the GD model under the ML framework. In the BR model, the four branches leading to the last common ancestor (LCA) of all *Prochlorococcus*, of the HL, LLI and LLII/III clades, of the HL and LLI clades, and of the HL clade, were allowed to have branch-specific turnover rates, whereas other branches were assumed to share the same rate. To avoid local optima, we ran 100 replicates for each ML analysis using different starting values (-start_val 1 accompanied with distinct seeds [-seed] provided by a random number generator). The likelihood of different runs among distinct models were compared (Fig. S7A). The presented estimates were based on the run with the maximum likelihood in each selected model.

#### *3.3 Gene gain and loss data integration*

For both methods, the corresponding results were compared and summarized to determine the common patterns shared by all analyses. AnGST categorizes the variation of genome contents into born, loss, duplication and horizontal gene transfer, whereas BadiRate only reports gene gain and loss through copy number changes. To smooth the comparison of all attempts, we standardized a "gain" event as the increase in the copy number of a gene family (including born, duplication and HGT), and accordingly a "loss" event as the decrease in the copy number of a gene family (including complete and partial loss) (Fig. S7C). Since the two methods gave a similar pattern of genome size reduction, we presented the number of gene gains and losses derived from the AnGST in the main text.

### 4. Calculating the rate of nonsynonymous nucleotide substitutions leading to radical and conservative amino acid changes, respectively

Previous study identified an excess of radical changes in *Prochlorococcus* HL and LLI/II/III lineages in comparison to their LLIV relatives (2). Here, radical changes are defined as nonsynonymous nucleotide substitutions leading to the replacements between amino acids with distinct physicochemical properties (charge, volume and polarity; Table S2), while conservative changes are among similar amino acids. The Radical and Conservative change Calculator (RCCalculator http://www.geneorder.org/RCCalculator/) was developed to compute the radical and conservative substitution rates ($d_R$ and $d_C$) which takes into account the GC biases of the DNA sequences (2).

In the present study, a total of 543 single-copy orthologous gene families, shared by all the 61 genomes of *Prochlorococcus* and *Synechococcus* clade 5.1/5.2, were retrieved from the aforementioned results of OrthoFinder v2.2.1 (66). Genes were aligned at the amino acid level using MAFFT v7.271(18), and DNA sequences were imposed on the alignments. Gaps and codons with ambiguous nucleotides were removed. The ratio of nonsynonoymous to synonymous substitution rates ($d_N/d_S$) was calculated using KaKs_Calculator under YN model for each of the orthologous gene pairs (73), and the median value of each gene family was used for RCCalculator. The transition/transversion ratio ($t_S/t_V$) of each gene family, also required by RCCalculator, was estimated using MEGA-CC v7.0.26 (74). By incorporating the uncultivated lineages of *Prochlorococcus*, a total of 751 single-copy orthologous gene families shared by 62 out of 65 genomes were retrieved and subject to the same procedures as described.

A total of six cases were considered for the calculation of $d_R$ and $d_C$, including two ways of categorizing amino acids (by charge and by volume and polarity) and three approaches of GC-bias correction (uncorrected, on codon frequency correction, and on amino acid composition correction). Under each case, given a gene family, RCCalculator estimates the number of radical and conservative sites for each sequence ($R_i$ and $C_i$, where $i \in [1, 61]$), as well as the numbers of radical and conservative differences of each sequence pair ($\Delta R_{ij}$ and $\Delta C_{ij}$, where $i \in [1, 61]$, $j \in [1, 61]$, and $i \neq j$). Then, the pairwise $d_R/d_C$ ratio was defined as $[\frac{\Delta R_{ij}}{mean(R_i,R_j)}] \Big/ [\frac{\Delta C_{ij}}{mean(C_i,C_j)}]$, where $i \in [1, 61]$, $j \in [1, 61]$, and $i \neq j$. In our study, each gene family had approximately 240 $d_R/d_C$ ratios resulted from the comparisons between sequences of the target group and the reference group (40 genomes in the target group vs. six genomes in the reference group), and 90

$d_R/d_C$ ratios from the control vs. reference comparisons (15 genomes in the control group vs. six reference ones). The mean values of these two categories were then used to represent the “target” and “control” $d_R/d_C$ ratios of the gene family. Further, after pooling all the 543 pairs of $d_R/d_C$ ratios together, sign test and paired t-test were used to determine significant differences between the $d_R/d_C$ ratios from the “target” and “control” groups.

**Fig. S1**

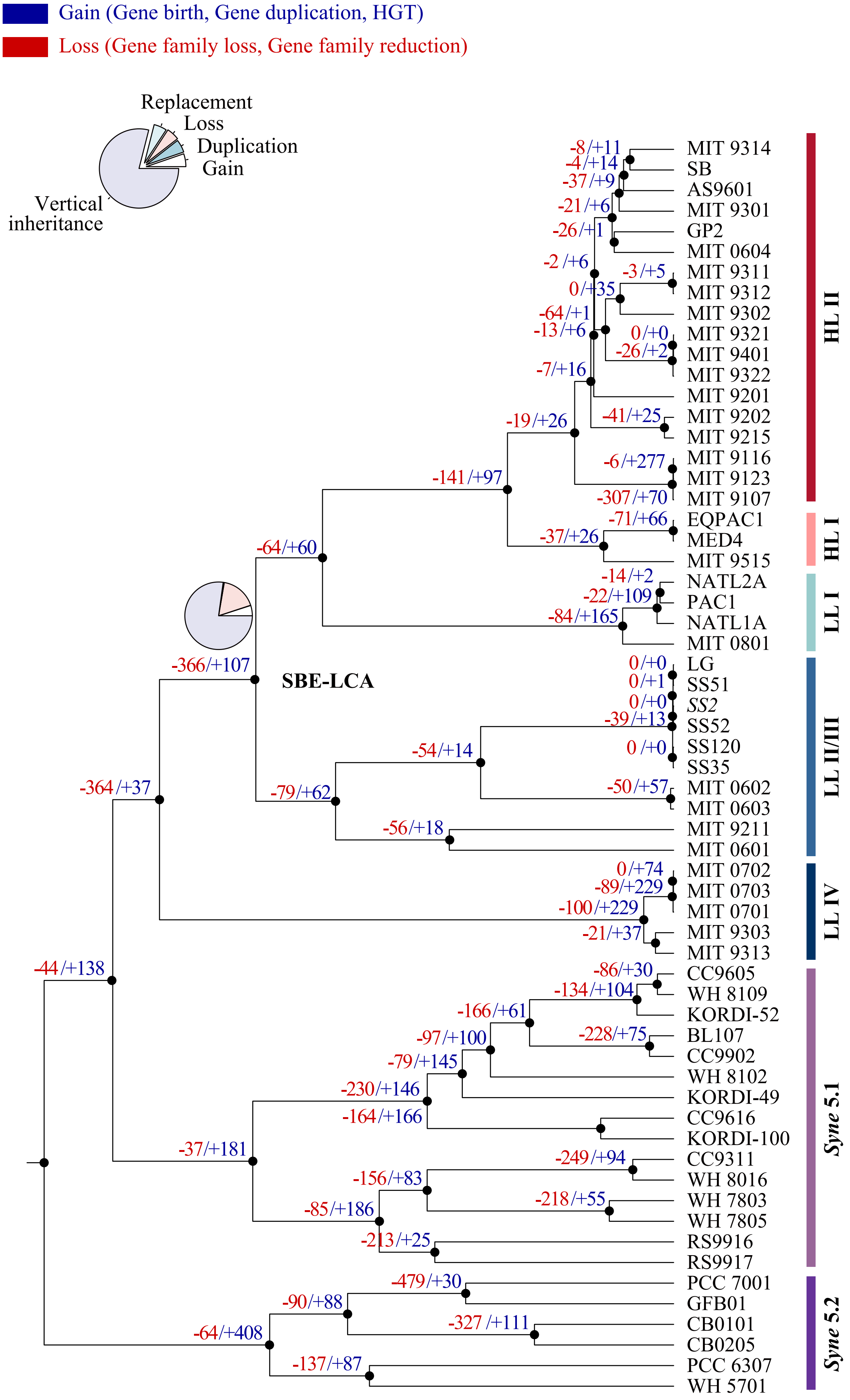


Fig. S1 The number of gene gain and loss events along the genome tree of *Prochlorococcus* and *Synechococcus* reconstructed by AnGST. Gene gain events include gene birth, duplication and HGT, while gene loss events comprise gene family size reduction and loss of entire gene families. The pie chart on the ancestral branches leading to SBE-LCA provides the detailed proportion of each type of genomic event in these key evolutionary stages.

**Fig. S2**

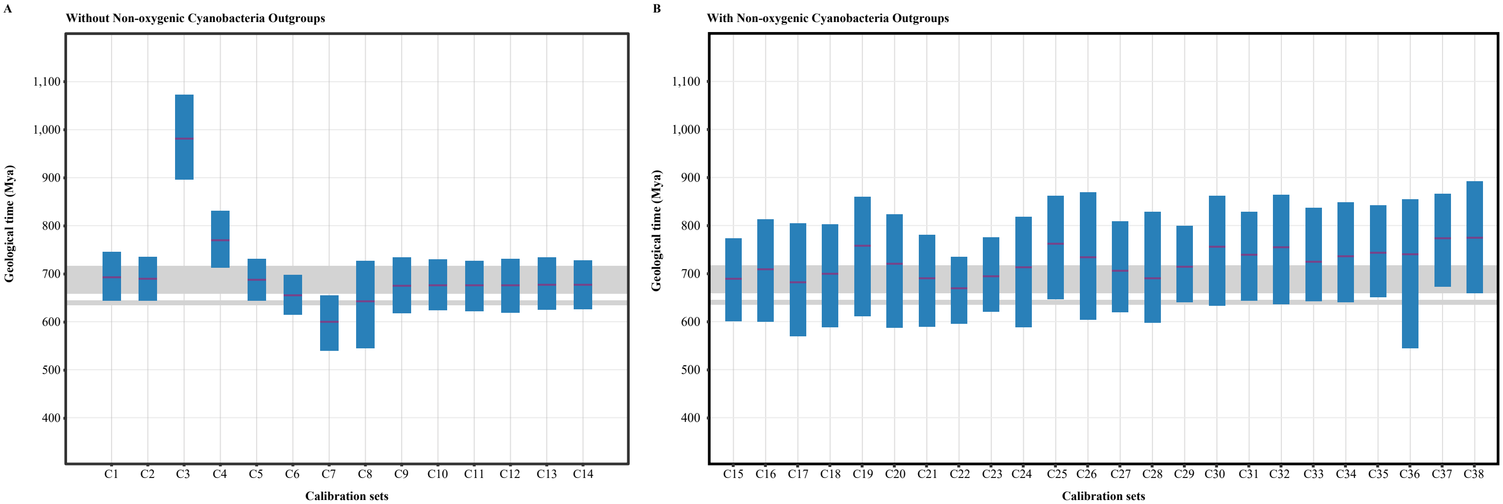


Fig. S2 Divergence time estimates of the ancestral node 'SBE-LCA' based on different calibration sets. (A) Calibration sets used for the phylgeony of oxygenic Cyanobacteria group, including some adapted from previous studies (C1-C6; Table S1) and others modified in the current study (C7-C14; Table S1). (B) Calibration sets used for the phylogeny of Cyanobacteria phylum including both oxygenic and non-oxygenic groups (C15-C38; Table S1). The purple lines and blue vertical bars represent the posterior age estimates and the 95% highest probability density (HPD) intervals, respectively. The upper and the lower horizontal grey bars represent the time of Sturtian glaciation and Marinoan glaciation, respectively.

Fig. S3

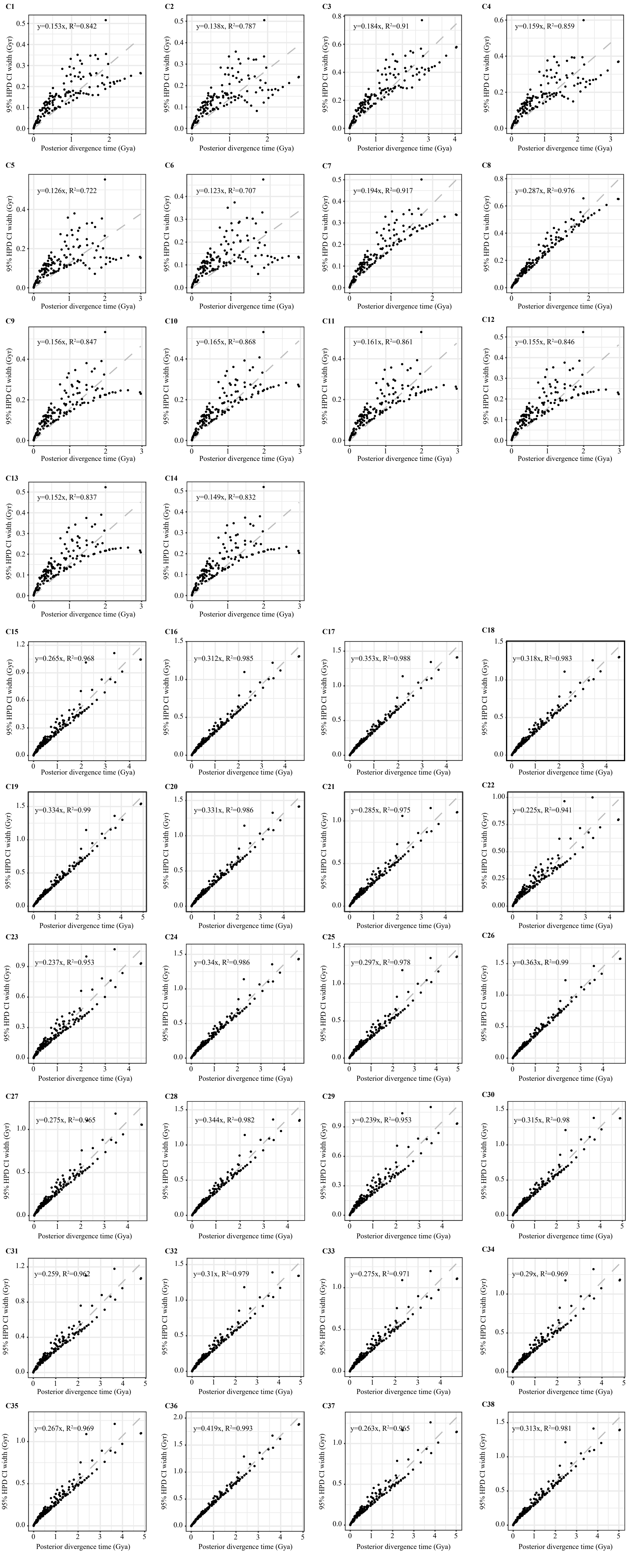


Fig. S3 The infinite-site plots of time estimates based on different calibration sets (C1-C38; see Table S1). The width of the 95% highest probability density (HPD) interval was plotted against the posterior means of the divergence time. A lower slope of the regression line suggests a higher precision of the molecular clock analysis.

Fig. S4 Phylogenomic trees of cyanobacteria based on concatenation of single-copy orthologous gene families at the amino acid sequence level. (A) Maximum likelihood phylogeny of 159 oxygenic Cyanobacteria genomes based on the 214 single-copy gene families shared by these genomes. (B) Maximum likelihood phylogeny of 159 oxygenic Cyanobacteria genomes based on the 90 gene families with evidence of compositional homogeneity in the protein sequences. (C) Bayesian inference phylogeny of 159 oxygenic Cyanobacteria genomes based on the 214 gene families. (D) Bayesian inference phylogeny of 159 oxygenic Cyanobacteria genomes based on the 90 gene families with evidence of compositional homogeneity in the protein sequences. (E) Maximum likelihood phylogeny of 159 oxygenic Cyanobacteria genomes as well as eight non-oxygenic Cyanobacteria outgroups based on the 90 gene families with evidence of compositional homogeneity in the protein sequences. Trees shown in (D) and (E) are used for molecular dating analyses, and calibrated ancestor nodes are marked with solid orange circle. The taxonomic classification on phylogeny is adapted from Sanchez-Baracaldo et al. (2015). Solid and open circles at ancestral nodes indicate the percentage of posterior probability or the frequency of the group defined by that node in 100 bootstrapped replicates is at least 95 and 85, respectively.

Macrocyanobacteria

Microcyanobacteria

Basal lineage

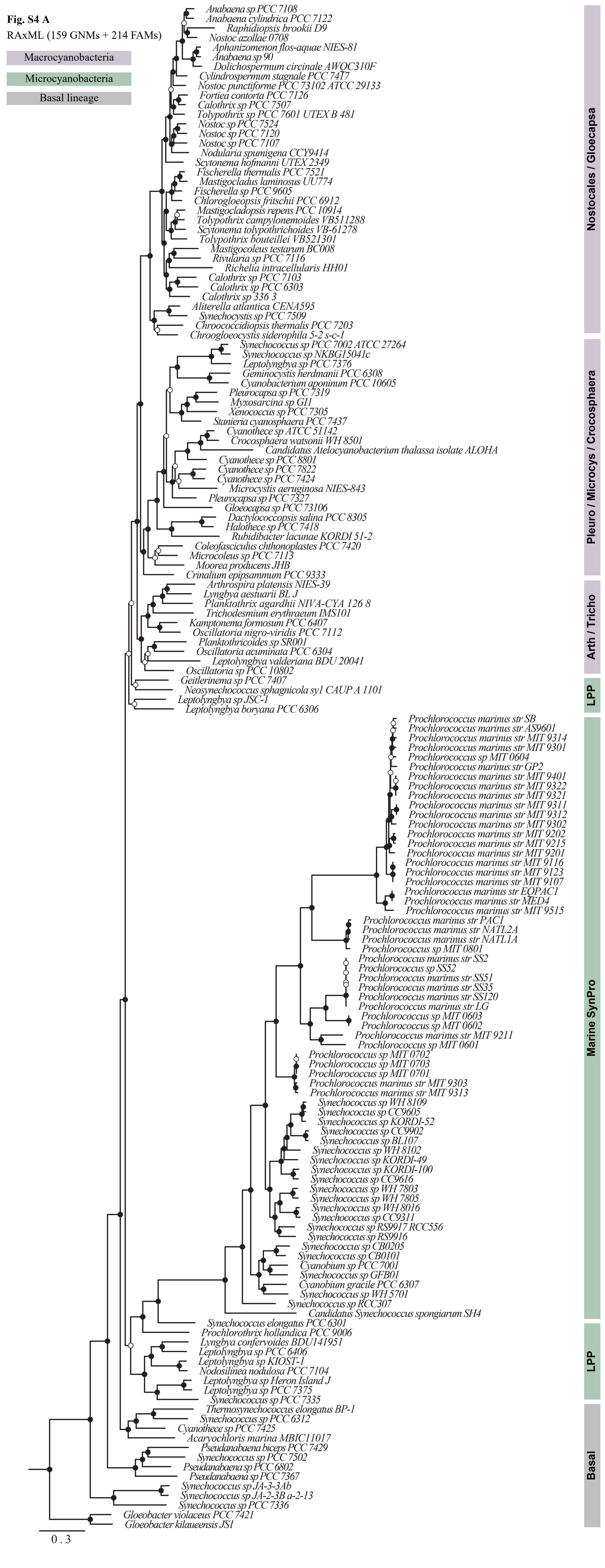
Fig. S4 A
RAxML (159 GNMs + 214 FAMs)
Macrocyanobacteria
Microcyanobacteria
Basal lineage
Anabaena sp PCC 7108
Anabaena cylindrica PCC 7122
Raphidiopsis brookii D9
Nostoc azollae 0708
Aphanizomenon flos-aquae NIES-81
Anabaena sp 90
Dolichospermum circinale AWQC310F
Cylindrospermum stagnale PCC 7417
Nostoc punctiforme PCC 73102 ATCC 29133
Fortiea contorta PCC 7126
Calothrix sp PCC 7507
Tolypothrix sp PCC 7601 UTEX B 481
Nostoc sp PCC 7524
Nostoc sp PCC 7120
Nostoc sp PCC 7107
Nodularia spumigena CCY9414
Scytonema hofmanni UTEX 2349
Fischerella thermalis PCC 7521
Mastigocladus laminosus UU774
Fischerella sp PCC 9605
Chlorogloeopsis fritschii PCC 6912
Mastigocladopsis repens PCC 10914
Tolypothrix campylonemoides VB511288
Scytonema tolypothrichoides VB-61278
Tolypothrix bouteillei VB521301
Mastigocoleus testarum BC008
Rivularia sp PCC 7116
Richelia intracellularis HH01
Calothrix sp PCC 7103
Calothrix sp PCC 6303
Calothrix sp 336 3
Aliterella atlantica CENA595
Synechocystis sp PCC 7509
Chroococcidiopsis thermalis PCC 7203
Chroogloeocystis siderophila 5-2 s-c-1
Synechococcus sp PCC 7002 ATCC 27264
Synechococcus sp NKBG15041c
Leptolyngbya sp PCC 7376
Geminocystis herdmanii PCC 6308
Cyanobacterium aponinum PCC 10605
Pleurocapsa sp PCC 7319
Myxosarcina sp GI1
Xenococcus sp PCC 7305
Stanieria cyanosphaera PCC 7437
Cyanothece sp ATCC 51142
Crocosphaera watsonii WH 8501
Candidatus Atelocyanobacterium thalassa isolate ALOHA
Cyanothece sp PCC 8801
Cyanothece sp PCC 7822
Cyanothece sp PCC 7424
Microcystis aeruginosa NIES-843
Pleurocapsa sp PCC 7327
Gloeocapsa sp PCC 73106
Dactylococcopsis salina PCC 8305
Halothece sp PCC 7418
Rubidibacter lacunae KORDI 51-2
Coleofasciculus chthonoplastes PCC 7420
Microcoleus sp PCC 7113
Moorea producens JHB
Crinalium epipsammum PCC 9333
Arthrospira platensis NIES-39
Lyngbya aestuarii BL J
Planktothrix agardhii NIVA-CYA 126 8
Trichodesmium erythraeum IMS101
Kamptonema formosum PCC 6407
Oscillatoria nigro-viridis PCC 7112
Planktothricoides sp SR001
Oscillatoria acuminata PCC 6304
Leptolyngbya valderiana BDU 20041
Oscillatoria sp PCC 10802
Geitlerinema sp PCC 7407
Neosynechococcus sphagnicola sy1 CAUP A 1101
Leptolyngbya sp JSC-1
Leptolyngbya boryana PCC 6306
Prochlorococcus marinus str SB
Prochlorococcus marinus str AS9601
Prochlorococcus marinus str MIT 9314
Prochlorococcus marinus str MIT 9301
Prochlorococcus sp MIT 0604
Prochlorococcus marinus str GP2
Prochlorococcus marinus str MIT 9401
Prochlorococcus marinus str MIT 9322
Prochlorococcus marinus str MIT 9321
Prochlorococcus marinus str MIT 9311
Prochlorococcus marinus str MIT 9312
Prochlorococcus marinus str MIT 9302
Prochlorococcus marinus str MIT 9202
Prochlorococcus marinus str MIT 9215
Prochlorococcus marinus str MIT 9201
Prochlorococcus marinus str MIT 9116
Prochlorococcus marinus str MIT 9123
Prochlorococcus marinus str MIT 9107
Prochlorococcus marinus str EQPAC1
Prochlorococcus marinus str MED4
Prochlorococcus marinus str MIT 9515
Prochlorococcus marinus str PAC1
Prochlorococcus marinus str NATL2A
Prochlorococcus marinus str NATL1A
Prochlorococcus sp MIT 0801
Prochlorococcus marinus str SS2
Prochlorococcus sp SS52
Prochlorococcus marinus str SS51
Prochlorococcus marinus str SS35
Prochlorococcus marinus str SS120
Prochlorococcus marinus str LG
Prochlorococcus sp MIT 0603
Prochlorococcus sp MIT 0602
Prochlorococcus marinus str MIT 9211
Prochlorococcus sp MIT 0601
Prochlorococcus sp MIT 0702
Prochlorococcus sp MIT 0703
Prochlorococcus sp MIT 0701
Prochlorococcus marinus str MIT 9303
Prochlorococcus marinus str MIT 9313
Synechococcus sp WH 8109
Synechococcus sp CC9605
Synechococcus sp KORDI-52
Synechococcus sp CC9902
Synechococcus sp BL107
Synechococcus sp WH 8102
Synechococcus sp KORDI-49
Synechococcus sp KORDI-100
Synechococcus sp CC9616
Synechococcus sp WH 7803
Synechococcus sp WH 7805
Synechococcus sp WH 8016
Synechococcus sp CC9311
Synechococcus sp RS9917 RCC556
Synechococcus sp RS9916
Synechococcus sp CB0205
Synechococcus sp CB0101
Cyanobium sp PCC 7001
Synechococcus sp GFB01
Cyanobium gracile PCC 6307
Synechococcus sp WH 5701
Synechococcus sp RCC307
Candidatus Synechococcus spongiarum SH4
Synechococcus elongatus PCC 6301
Prochlorothrix hollandica PCC 9006
Lyngbya confervoides BDU141951
Leptolyngbya sp PCC 6406
Leptolyngbya sp KIOST-1
Nodosilinea nodulosa PCC 7104
Leptolyngbya sp Heron Island J
Leptolyngbya sp PCC 7375
Synechococcus sp PCC 7335
Thermosynechococcus elongatus BP-1
Synechococcus sp PCC 6312
Cyanothece sp PCC 7425
Acaryochloris marina MBIC11017
Pseudanabaena biceps PCC 7429
Synechococcus sp PCC 7502
Pseudanabaena sp PCC 6802
Pseudanabaena sp PCC 7367
Synechococcus sp JA-3-3Ab
Synechococcus sp JA-2-3B a-2-13
Synechococcus sp PCC 7336
Gloeobacter violaceus PCC 7421
Gloeobacter kilaueensis JS1
0 . 3
Nostocales / Gloecapsa
Pleuro / Microcys / Crocosphaera
Arth / Tricho
LPP
Marine SynPro
LPP
Basal

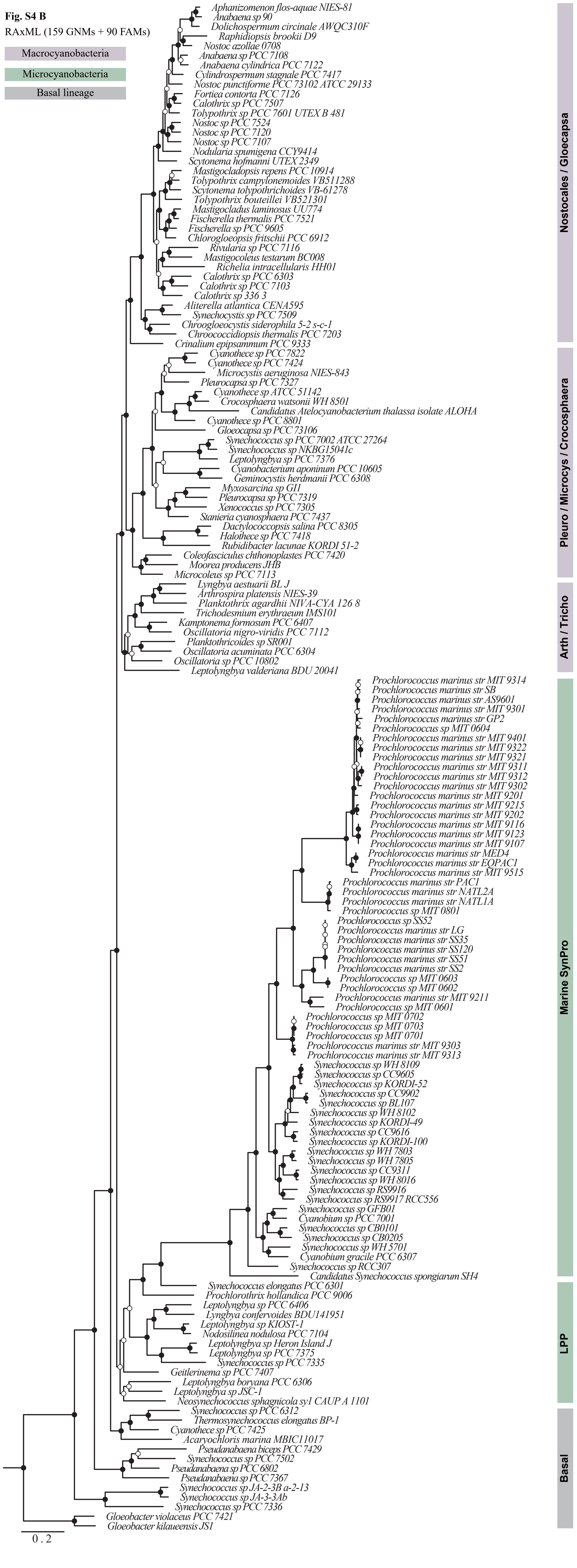


**Fig. S4 B**

RAxML (159 GNMs + 90 FAMs)

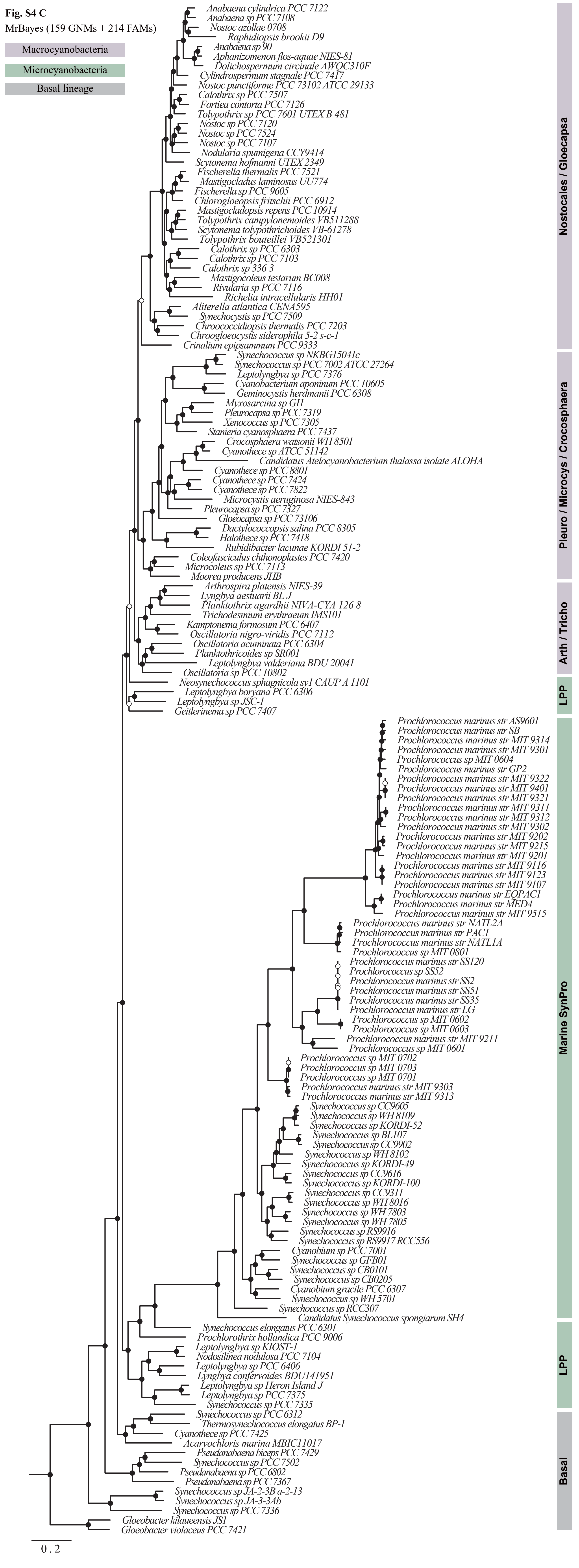
Fig. S4 C
MrBayes (159 GNMs + 214 FAMs)
Macrocyanobacteria
Microcyanobacteria
Basal lineage
Anabaena cylindrica PCC 7122
Anabaena sp PCC 7108
Nostoc azollae 0708
Raphidiopsis brookii D9
Anabaena sp 90
Aphanizomenon flos-aquae NIES-81
Dolichospermum circinale AWQC310F
Cylindrospermum stagnale PCC 7417
Nostoc punctiforme PCC 73102 ATCC 29133
Calothrix sp PCC 7507
Fortiea contorta PCC 7126
Tolypothrix sp PCC 7601 UTEX B 481
Nostoc sp PCC 7120
Nostoc sp PCC 7524
Nostoc sp PCC 7107
Nodularia spumigena CCY9414
Scytonema hofmanni UTEX 2349
Fischerella thermalis PCC 7521
Mastigocladus laminosus UU774
Fischerella sp PCC 9605
Chlorogloeopsis fritschii PCC 6912
Mastigocladopsis repens PCC 10914
Tolypothrix campylonemoides VB511288
Scytonema tolypothrichoides VB-61278
Tolypothrix bouteillei VB521301
Calothrix sp PCC 6303
Calothrix sp PCC 7103
Calothrix sp 336 3
Mastigocoleus testarum BC008
Rivularia sp PCC 7116
Richelia intracellularis HH01
Aliterella atlantica CENA595
Synechocystis sp PCC 7509
Chroococcidiopsis thermalis PCC 7203
Chroogloeocystis siderophila 5-2 s-c-1
Crinalium epipsammum PCC 9333
Synechococcus sp NKBG15041c
Synechococcus sp PCC 7002 ATCC 27264
Leptolyngbya sp PCC 7376
Cyanobacterium aponinum PCC 10605
Geminocystis herdmanii PCC 6308
Myxosarcina sp GI1
Pleurocapsa sp PCC 7319
Xenococcus sp PCC 7305
Stanieria cyanosphaera PCC 7437
Crocosphaera watsonii WH 8501
Cyanothece sp ATCC 51142
Candidatus Atelocyanobacterium thalassa isolate ALOHA
Cyanothece sp PCC 8801
Cyanothece sp PCC 7424
Cyanothece sp PCC 7822
Microcystis aeruginosa NIES-843
Pleurocapsa sp PCC 7327
Gloeocapsa sp PCC 73106
Dactylococcopsis salina PCC 8305
Halothece sp PCC 7418
Rubidibacter lacunae KORDI 51-2
Coleofasciculus chthonoplastes PCC 7420
Microcoleus sp PCC 7113
Moorea producens JHB
Arthrospira platensis NIES-39
Lyngbya aestuarii BL J
Planktothrix agardhii NIVA-CYA 126 8
Trichodesmium erythraeum IMS101
Kamptonema formosum PCC 6407
Oscillatoria nigro-viridis PCC 7112
Oscillatoria acuminata PCC 6304
Planktothricoides sp SR001
Leptolyngbya valderiana BDU 20041
Oscillatoria sp PCC 10802
Neosynechococcus sphagnicola sy1 CAUP A 1101
Leptolyngbya boryana PCC 6306
Leptolyngbya sp JSC-1
Geitlerinema sp PCC 7407
Prochlorococcus marinus str AS9601
Prochlorococcus marinus str SB
Prochlorococcus marinus str MIT 9314
Prochlorococcus marinus str MIT 9301
Prochlorococcus sp MIT 0604
Prochlorococcus marinus str GP2
Prochlorococcus marinus str MIT 9322
Prochlorococcus marinus str MIT 9401
Prochlorococcus marinus str MIT 9321
Prochlorococcus marinus str MIT 9311
Prochlorococcus marinus str MIT 9312
Prochlorococcus marinus str MIT 9302
Prochlorococcus marinus str MIT 9202
Prochlorococcus marinus str MIT 9215
Prochlorococcus marinus str MIT 9201
Prochlorococcus marinus str MIT 9116
Prochlorococcus marinus str MIT 9123
Prochlorococcus marinus str MIT 9107
Prochlorococcus marinus str EQPAC1
Prochlorococcus marinus str MED4
Prochlorococcus marinus str MIT 9515
Prochlorococcus marinus str NATL2A
Prochlorococcus marinus str PAC1
Prochlorococcus marinus str NATL1A
Prochlorococcus sp MIT 0801
Prochlorococcus marinus str SS120
Prochlorococcus sp SS52
Prochlorococcus marinus str SS2
Prochlorococcus marinus str SS51
Prochlorococcus marinus str SS35
Prochlorococcus marinus str LG
Prochlorococcus sp MIT 0602
Prochlorococcus sp MIT 0603
Prochlorococcus marinus str MIT 9211
Prochlorococcus sp MIT 0601
Prochlorococcus sp MIT 0702
Prochlorococcus sp MIT 0703
Prochlorococcus sp MIT 0701
Prochlorococcus marinus str MIT 9303
Prochlorococcus marinus str MIT 9313
Synechococcus sp CC9605
Synechococcus sp WH 8109
Synechococcus sp KORDI-52
Synechococcus sp BL107
Synechococcus sp CC9902
Synechococcus sp WH 8102
Synechococcus sp KORDI-49
Synechococcus sp CC9616
Synechococcus sp KORDI-100
Synechococcus sp CC9311
Synechococcus sp WH 8016
Synechococcus sp WH 7803
Synechococcus sp WH 7805
Synechococcus sp RS9916
Synechococcus sp RS9917 RCC556
Cyanobium sp PCC 7001
Synechococcus sp GFB01
Synechococcus sp CB0101
Synechococcus sp CB0205
Cyanobium gracile PCC 6307
Synechococcus sp WH 5701
Synechococcus sp RCC307
Candidatus Synechococcus spongiarum SH4
Synechococcus elongatus PCC 6301
Prochlorothrix hollandica PCC 9006
Leptolyngbya sp KIOST-1
Nodosilinea nodulosa PCC 7104
Leptolyngbya sp PCC 6406
Lyngbya confervoides BDU141951
Leptolyngbya sp Heron Island J
Leptolyngbya sp PCC 7375
Synechococcus sp PCC 7335
Synechococcus sp PCC 6312
Thermosynechococcus elongatus BP-1
Cyanothece sp PCC 7425
Acaryochloris marina MBIC11017
Pseudanabaena biceps PCC 7429
Synechococcus sp PCC 7502
Pseudanabaena sp PCC 6802
Pseudanabaena sp PCC 7367
Synechococcus sp JA-2-3B a-2-13
Synechococcus sp JA-3-3Ab
Synechococcus sp PCC 7336
Gloeobacter kilaueensis JS1
Gloeobacter violaceus PCC 7421
0.2
Nostocales / Gloecapsa
Pleuro / Microcys / Crocosphaera
Arth / Tricho
LPP
Marine SynPro
LPP
Basal

**Fig. S4 D**

MrBayes (159 GNMs + 90 FAMs)

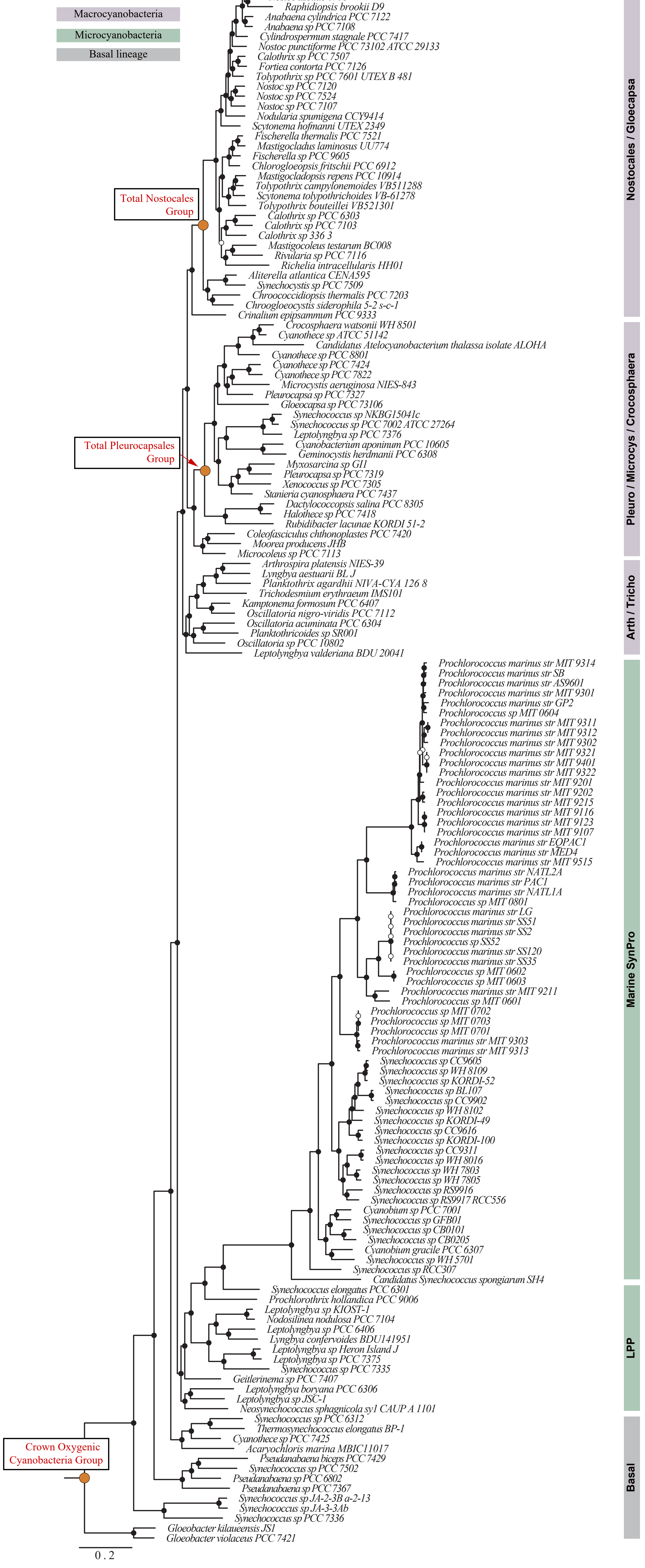

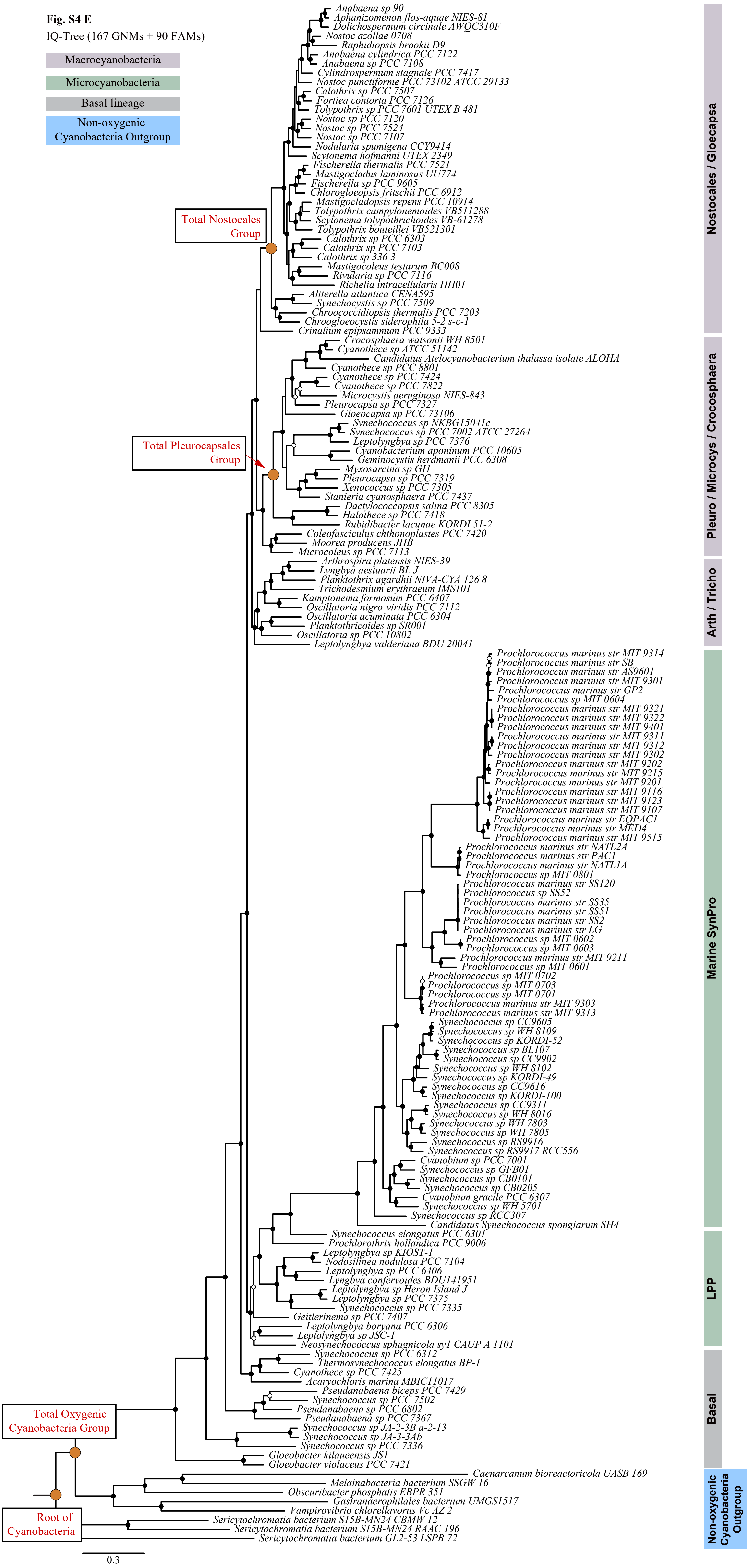
Fig. S4 E
IQ-Tree (167 GNMs + 90 FAMs)
Macrocyanobacteria
Microcyanobacteria
Basal lineage
Non-oxygenic Cyanobacteria Outgroup
Total Nostocales Group
Total Pleurocapsales Group
Total Oxygenic Cyanobacteria Group
Root of Cyanobacteria
Anabaena sp 90
Aphanizomenon flos-aquae NIES-81
Dolichospermum circinale AWQC310F
Nostoc azollae 0708
Raphidiopsis brookii D9
Anabaena cylindrica PCC 7122
Anabaena sp PCC 7108
Cylindrospermum stagnale PCC 7417
Nostoc punctiforme PCC 73102 ATCC 29133
Calothrix sp PCC 7507
Fortiea contorta PCC 7126
Tolypothrix sp PCC 7601 UTEX B 481
Nostoc sp PCC 7120
Nostoc sp PCC 7524
Nostoc sp PCC 7107
Nodularia spumigena CCY9414
Scytonema hofmanni UTEX 2349
Fischerella thermalis PCC 7521
Mastigocladus laminosus UU774
Fischerella sp PCC 9605
Chlorogloeopsis fritschii PCC 6912
Mastigocladopsis repens PCC 10914
Tolypothrix campylonemoides VB511288
Scytonema tolypothrichoides VB-61278
Tolypothrix bouteillei VB521301
Calothrix sp PCC 6303
Calothrix sp PCC 7103
Calothrix sp 336 3
Mastigocoleus testarum BC008
Rivularia sp PCC 7116
Richelia intracellularis HH01
Aliterella atlantica CENA595
Synechocystis sp PCC 7509
Chroococcidiopsis thermalis PCC 7203
Chroogloeocystis siderophila 5-2 s-c-1
Crinalium epipsammum PCC 9333
Crocosphaera watsonii WH 8501
Cyanothece sp ATCC 51142
Candidatus Atelocyanobacterium thalassa isolate ALOHA
Cyanothece sp PCC 8801
Cyanothece sp PCC 7424
Cyanothece sp PCC 7822
Microcystis aeruginosa NIES-843
Pleurocapsa sp PCC 7327
Gloeocapsa sp PCC 73106
Synechococcus sp NKBG15041c
Synechococcus sp PCC 7002 ATCC 27264
Leptolyngbya sp PCC 7376
Cyanobacterium aponinum PCC 10605
Geminocystis herdmanii PCC 6308
Myxosarcina sp GI1
Pleurocapsa sp PCC 7319
Xenococcus sp PCC 7305
Stanieria cyanosphaera PCC 7437
Dactylococcopsis salina PCC 8305
Halothece sp PCC 7418
Rubidibacter lacunae KORDI 51-2
Coleofasciculus chthonoplastes PCC 7420
Moorea producens JHB
Microcoleus sp PCC 7113
Arthrospira platensis NIES-39
Lyngbya aestuarii BL J
Planktothrix agardhii NIVA-CYA 126 8
Trichodesmium erythraeum IMS101
Kamptonema formosum PCC 6407
Oscillatoria nigro-viridis PCC 7112
Oscillatoria acuminata PCC 6304
Planktothricoides sp SR001
Oscillatoria sp PCC 10802
Leptolyngbya valderiana BDU 20041
Prochlorococcus marinus str MIT 9314
Prochlorococcus marinus str SB
Prochlorococcus marinus str AS9601
Prochlorococcus marinus str MIT 9301
Prochlorococcus marinus str GP2
Prochlorococcus sp MIT 0604
Prochlorococcus marinus str MIT 9321
Prochlorococcus marinus str MIT 9322
Prochlorococcus marinus str MIT 9401
Prochlorococcus marinus str MIT 9311
Prochlorococcus marinus str MIT 9312
Prochlorococcus marinus str MIT 9302
Prochlorococcus marinus str MIT 9202
Prochlorococcus marinus str MIT 9215
Prochlorococcus marinus str MIT 9201
Prochlorococcus marinus str MIT 9116
Prochlorococcus marinus str MIT 9123
Prochlorococcus marinus str MIT 9107
Prochlorococcus marinus str EQPAC1
Prochlorococcus marinus str MED4
Prochlorococcus marinus str MIT 9515
Prochlorococcus marinus str NATL2A
Prochlorococcus marinus str PAC1
Prochlorococcus marinus str NATL1A
Prochlorococcus sp MIT 0801
Prochlorococcus marinus str SS120
Prochlorococcus sp SS52
Prochlorococcus marinus str SS35
Prochlorococcus marinus str SS51
Prochlorococcus marinus str SS2
Prochlorococcus marinus str LG
Prochlorococcus sp MIT 0602
Prochlorococcus sp MIT 0603
Prochlorococcus marinus str MIT 9211
Prochlorococcus sp MIT 0601
Prochlorococcus sp MIT 0702
Prochlorococcus sp MIT 0703
Prochlorococcus sp MIT 0701
Prochlorococcus marinus str MIT 9303
Prochlorococcus marinus str MIT 9313
Synechococcus sp CC9605
Synechococcus sp WH 8109
Synechococcus sp KORDI-52
Synechococcus sp BL107
Synechococcus sp CC9902
Synechococcus sp WH 8102
Synechococcus sp KORDI-49
Synechococcus sp CC9616
Synechococcus sp KORDI-100
Synechococcus sp CC9311
Synechococcus sp WH 8016
Synechococcus sp WH 7803
Synechococcus sp WH 7805
Synechococcus sp RS9916
Synechococcus sp RS9917 RCC556
Cyanobium sp PCC 7001
Synechococcus sp GFB01
Synechococcus sp CB0101
Synechococcus sp CB0205
Cyanobium gracile PCC 6307
Synechococcus sp WH 5701
Synechococcus sp RCC307
Candidatus Synechococcus spongiarum SH4
Synechococcus elongatus PCC 6301
Prochlorothrix hollandica PCC 9006
Leptolyngbya sp KIOST-1
Nodosilinea nodulosa PCC 7104
Leptolyngbya sp PCC 6406
Lyngbya confervoides BDU141951
Leptolyngbya sp Heron Island J
Leptolyngbya sp PCC 7375
Synechococcus sp PCC 7335
Geitlerinema sp PCC 7407
Leptolyngbya boryana PCC 6306
Leptolyngbya sp JSC-1
Neosynechococcus sphagnicola sy1 CAUP A 1101
Synechococcus sp PCC 6312
Thermosynechococcus elongatus BP-1
Cyanothece sp PCC 7425
Acaryochloris marina MBIC11017
Pseudanabaena biceps PCC 7429
Synechococcus sp PCC 7502
Pseudanabaena sp PCC 6802
Pseudanabaena sp PCC 7367
Synechococcus sp JA-2-3B a-2-13
Synechococcus sp JA-3-3Ab
Synechococcus sp PCC 7336
Gloeobacter kilaueensis JS1
Gloeobacter violaceus PCC 7421
Caenarcanum bioreactoricola UASB 169
Melainabacteria bacterium SSGW 16
Obscuribacter phosphatis EBPR 351
Gastranaerophilales bacterium UMGS1517
Vampirovibrio chlorellavorus Vc AZ 2
Sericytochromatia bacterium S15B-MN24 CBMW 12
Sericytochromatia bacterium S15B-MN24 RAAC 196
Sericytochromatia bacterium GL2-53 LSPB 72
0.3
Nostocales / Gloecapsa
Pleuro / Microcys / Crocosphaera
Arth / Tricho
Marine SynPro
LPP
Basal
Non-oxygenic Cyanobacteria Outgroup

**Fig. S5**

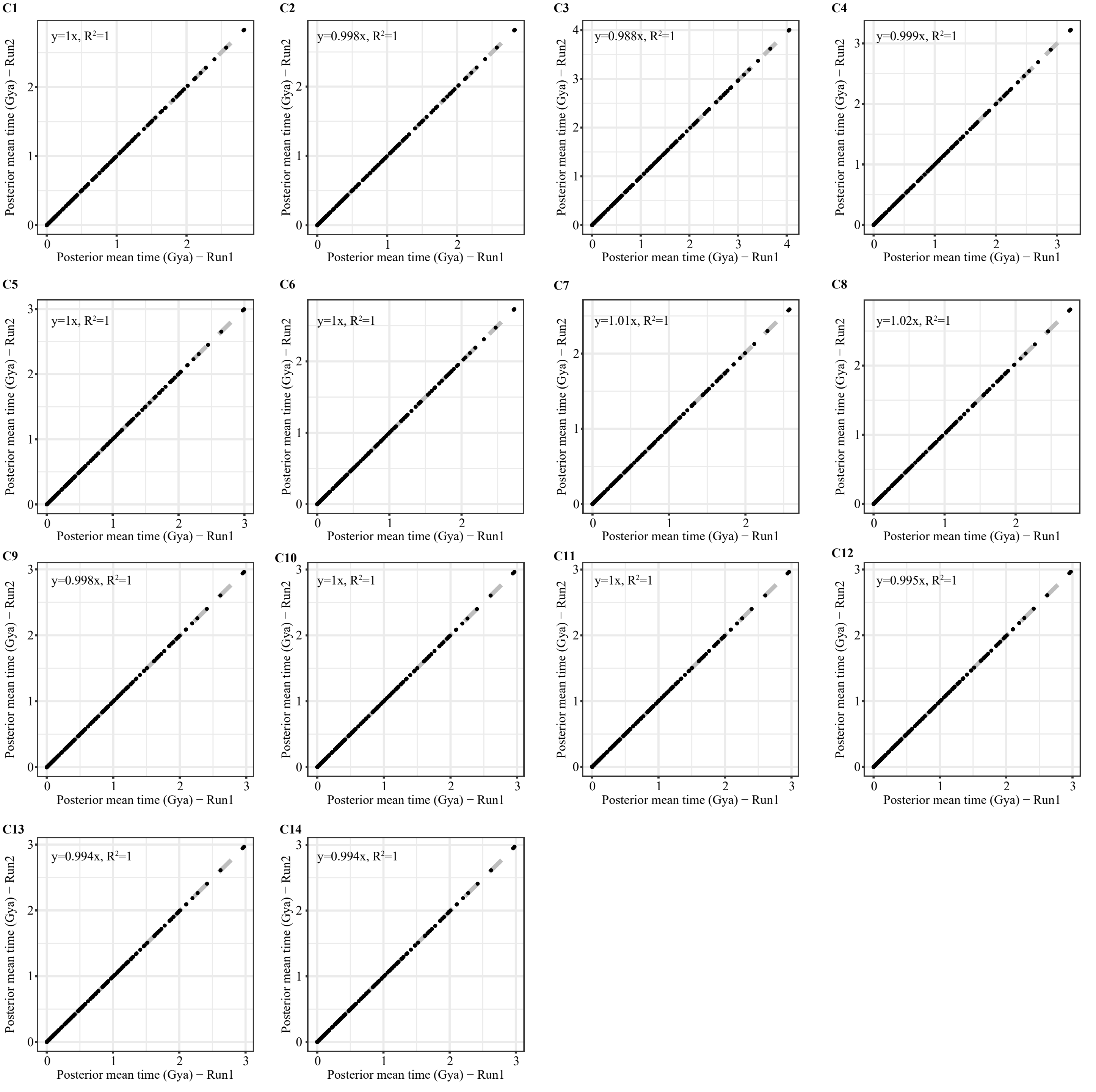


Fig. S5 Correlation of the posterior mean ages between replicated MCMC runs. Convergence of the independent run is achieved if points fall almost perfectly on the y=x line. The plots shown here are based on the calibration set C1-C14 (Table S1). The converged analyses based on the calibration set C15-C38 (Table S1) are not shown.

Fig. S6

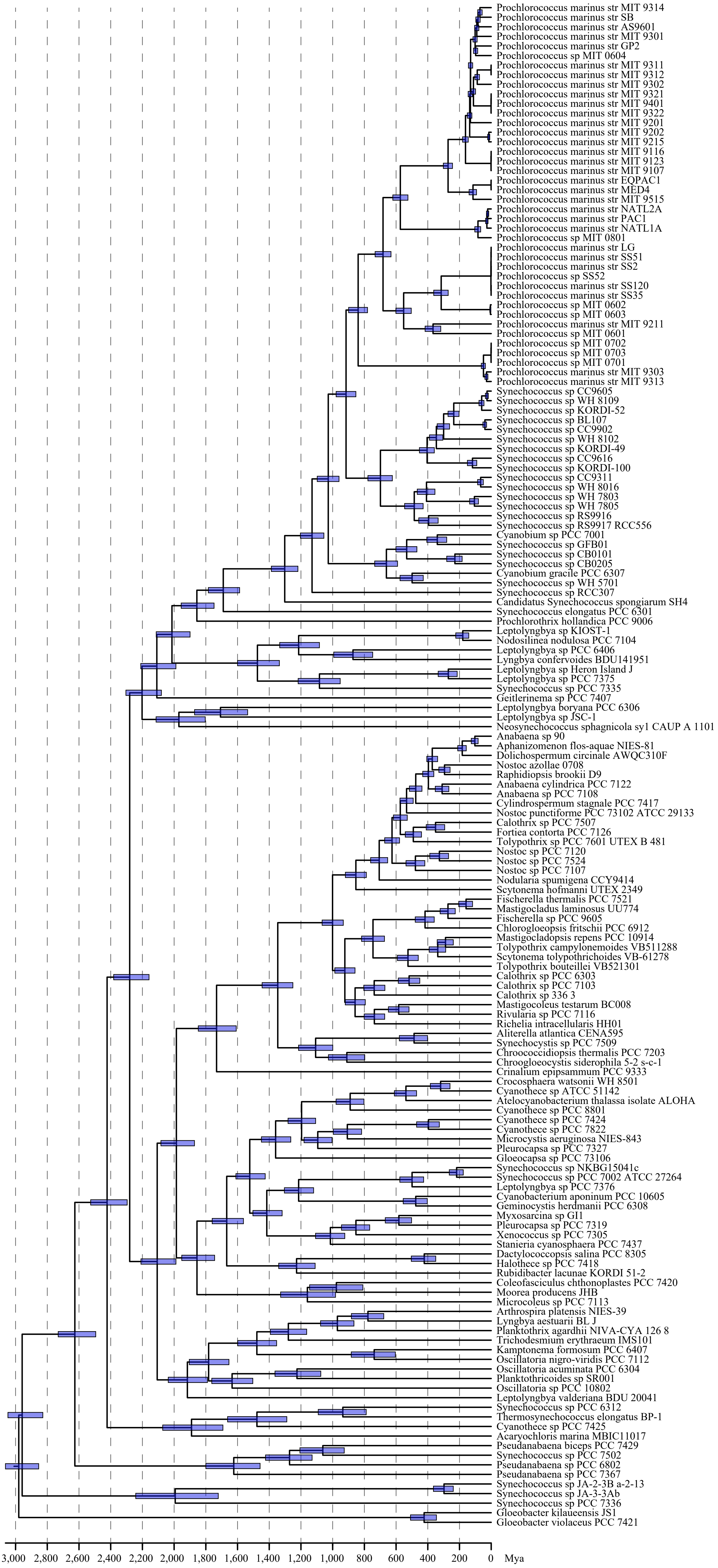


Fig. S6 A chronogram of cyanobacteria reconstructed with a relaxed molecular analysis implemented in MCMCTree. The molecular dating analysis uses 27 genes, a Bayesian phylogenomic tree of 159 genomes constructed with protein sequences of 90 gene families under the calibration set C14.

**Fig. S7**

A

| NAME | MEAN | MEDIAN | MAX |
|---|---|---|---|
| BDI+GR+ML | -111264.43 | -88323.13 | -84508.82 |
| GD+GR+ML | -85685.10 | -84551.35 | -84551.35 |
| LI+GR+ML | -97116.23 | -85956.71 | -85956.71 |
| GD+BR+ML | -100939.50 | -94705.62 | -91345.30 |
| BDI+BR+ML | -155458.90 | -107205.09 | -94802.30 |
| LI+BR+ML | -149392.61 | -115007.38 | -100625.35 |
| L+BR+ML | -152360.18 | -151987.12 | -128020.33 |
| L+GR+ML | -128738.83 | -128738.83 | -128738.83 |
| GD+FR+ML | -232431.80 | -230572.17 | -219759.66 |

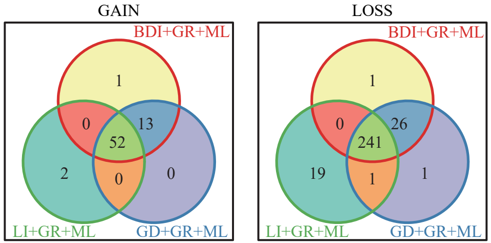


B

| Full Label | Description |
|---|---|
| MB214_C1 | HGT time consistency + Ultrametric tree derived from MCMCTREE + MrBayes (214 FAMs) + calibration set C1) |
| MB214_C7 | HGT time consistency + Ultrametric tree derived from MCMCTREE + MrBayes (114 FAMs) + calibration set C7) |
| MB90_C1 | HGT time consistency + Ultrametric tree derived from MCMCTREE + MrBayes (90 FAMs) + calibration set C1) |
| MB90_C7 | HGT time consistency + Ultrametric tree derived from MCMCTREE + MrBayes (90 FAMs) + calibration set C7) |

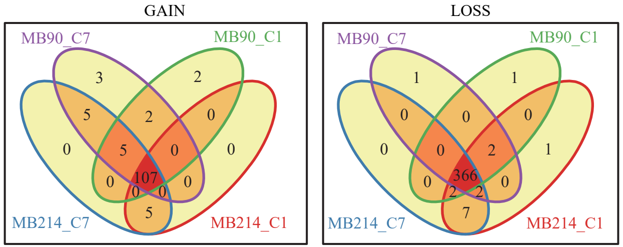


C

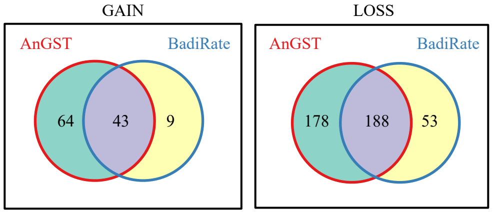


Fig. S7 (A) (Left) Multiple models are implemented in BadiRate for ancestral reconstruction, with the mean, median, and maximum likelihood values of each in 100 replicated analyses are shown. Models with their maximum likelihood values ranking at the top three (shaded in grey) are subject to further analyses. (Right) Venn diagrams show the number of gain and loss events, respectively, reconstructed with the three models shown in (Left). (B) (Left) For ancestral reconstructions with AnGST, a chronogram is used to limit HGT events occurring between contemporaneous lineages (HGT Time Consistency). Chronograms are estimated with MCMCTree based on 214 or 90 gene families (FAMs) under calibration set C1 or C7. (Right) Venn diagrams show the number of gain and loss events, respectively, reconstructed based on different strategies shown in (Left). (C) Venn diagrams show the number of gene families predicted to be gained or lost by AnGST and BadiRate during the evolutionary. The former was presented in the main text.

**Table S1** A list of fossil calibration sets employed in the present study. Maximum and minimum time constraints are in the unit of billion years ago (Ga). References for calibrations are provided.

| | Calibration Set | Cyanobacteria Root | Total Oxygenic Cyanobacteria | Crown Oxygenic Cyanobacteria | Total Pleurocapsales | Total Nostocales | Crown Nostocales |
|---|---|---|---|---|---|---|---|
| Without Non-oxygenic Cyanobacteria Outgroup | C1 [15] | - | - | 2.32-2.7 [1,2] | 1.7-2.45 [3,4] | - | 2.1-2.45 [3,5] |
| | C2 [16] | - | - | 2.32-2.7 | 1.7-1.9 [6,7,9] | - | 1.6-1.9 [6,7,8] |
| | C3 [15] | - | - | 2.32-3.0 [2,10,11,12] | 1.7-2.45 | - | 2.1-2.45 |
| | C4 [16] | - | - | 2.32-3.0 | 1.7-1.9 | - | 1.6-1.9 |
| | C5 [17] | - | - | 2.32-3.0 | 1.7-1.9 | - | <2.1 [5] |
| | C6 [17] | - | - | 2.32-2.7 | 1.7-1.9 | - | <2.1 |
| | C7 | - | - | 2.32-2.7 | - | - | - |
| | C8 | - | - | 2.32-3.0 | - | - | - |
| | C9 | - | - | 2.32-3.0 | >1.7 | >1.6 | - |
| | C10 | - | - | 2.32-3.0 | >1.7 | >1.9 | - |
| | C11 | - | - | 2.32-3.0 | >1.7 | >2.1 | - |
| | C12 | - | - | 2.32-3.0 | >1.9 | >1.6 | - |
| | C13 | - | - | 2.32-3.0 | >1.9 | >1.9 | - |
| | C14 | - | - | 2.32-3.0 | >1.9 | >2.1 | - |
| With Non-oxygenic Cyanobacteria Outgroup | C15 | <3.8 [13,14] | >3.0 [10,11,12] | - | >1.7 | >1.6 | - |
| | C16 | <3.8 | >3.0 | - | >1.7 | >1.9 | - |
| | C17 | <3.8 | >3.0 | - | >1.7 | >2.1 | - |
| | C18 | <3.8 | >3.0 | - | >1.9 | >1.6 | - |
| | C19 | <3.8 | >3.0 | - | >1.9 | >1.9 | - |
| | C20 | <3.8 | >3.0 | - | >1.9 | >2.1 | - |
| | C21 | <4.0 [13,14] | >3.0 | - | >1.7 | >1.6 | - |
| | C22 | <4.0 | >3.0 | - | >1.7 | >1.9 | - |
| | C23 | <4.0 | >3.0 | - | >1.7 | >2.1 | - |
| | C24 | <4.0 | >3.0 | - | >1.9 | >1.6 | - |
| | C25 | <4.0 | >3.0 | - | >1.9 | >1.9 | - |
| | C26 | <4.0 | >3.0 | - | >1.9 | >2.1 | - |
| | C27 | <4.2 [13,14] | >3.0 | - | >1.7 | >1.6 | - |
| | C28 | <4.2 | >3.0 | - | >1.7 | >1.9 | - |
| | C29 | <4.2 | >3.0 | - | >1.7 | >2.1 | - |
| | C30 | <4.2 | >3.0 | - | >1.9 | >1.6 | - |
| | C31 | <4.2 | >3.0 | - | >1.9 | >1.9 | - |
| | C32 | <4.2 | >3.0 | - | >1.9 | >2.1 | - |
| | C33 | <4.5 [13,14] | >3.0 | - | >1.7 | >1.6 | - |
| | C34 | <4.5 | >3.0 | - | >1.7 | >1.9 | - |
| | C35 | <4.5 | >3.0 | - | >1.7 | >2.1 | - |
| | C36 | <4.5 | >3.0 | - | >1.9 | >1.6 | - |
| | C37 | <4.5 | >3.0 | - | >1.9 | >1.9 | - |
| | C38 | <4.5 | >3.0 | - | >1.9 | >2.1 | - |

Table S2 Classification of amino acids by two independent schemes based on physiochemical properties of the amino acids.

| |
|---|
| Classification by charge (Hughes et al., 1990) |
| Positive R, H, K |
| Negative D, E |
| Neutral A, N, C, Q, G, I, L, M, F, P, S, T, W, Y, V |
| Classification by volume and polarity (Miyata et al., 1979) |
| Special C |
| Neutral and small A, G, P, S, T |
| Polar and relative small N, Q, D, E |
| Polar and relative large R, H, K |
| Nonpolar and relatively small I, L, M, V |
| Nonpolar and relatively large F, W, Y |

Hughes, A.L., Ota, T., and Nei, M. (1990) Positive Darwinian selection promotes charge profile diversity in the antigen-binding cleft of class I major-histocompatibility-complex molecules. Molecular biology and evolution 7: 515-524.

Miyata, T., Miyazawa, S., and Yasunaga, T. (1979) Two types of amino acid substitutions in protein evolution. Journal of Molecular Evolution 12: 219-236.
